# Metabolically-Incorporated Deuterium in Myelin Localized by Neutron Diffraction and Identified by Mass Spectrometry

**DOI:** 10.1101/2021.12.17.473245

**Authors:** Anne Baumann, Andrew R. Denninger, Marek Domin, Bruno Demé, Daniel A. Kirschner

**Affiliations:** Department of Biology, Boston College, Chestnut Hill, Massachusetts, United States of America; Division of Psychiatry, Haukeland University Hospital, Bergen, Norway; Department of Biomedicine, University of Bergen, Bergen, Norway; Department of Chemistry, Boston College, Chestnut Hill, Massachusetts, United States of America; Institut Laue-Langevin, Grenoble, France

## Abstract

Myelin is a natural and dynamic multilamellar membrane structure that continues to be of significant biological and neurological interest, especially with respect to its biosynthesis and assembly during its normal formation, maintenance, and pathological breakdown. To explore the usefulness of neutron diffraction in the structural analysis of myelin, we investigated the use of *in vivo* labeling by metabolically incorporating non-toxic levels of deuterium (^2^H; D) via drinking water into a pregnant dam (D-dam) and her developing embryos. All of the mice were sacrificed when the pups (D-pups) were 55 days old. Myelinated sciatic nerves were dissected, fixed in glutaraldehyde and examined by neutron diffraction. Parallel samples that were unfixed (trigeminal nerves) were frozen for mass spectrometry (MS). The diffraction patterns of the nerves from deuterium-fed mice (D-mice) vs. the controls (H-mice) had major differences in the intensities of the Bragg peaks but no appreciable differences in myelin periodicity. Neutron scattering density profiles showed an appreciable increase in density at the center of the lipid-rich membrane bilayer. This increase was greater in D-pups than in D-dam, and its localization was consistent with deuteration of lipid hydrocarbon, which predominates over transmembrane protein in myelin. MS analysis of the lipids isolated from the trigeminal nerves demonstrated that in the pups the percentage of lipids that had one or more deuterium atoms was uniformly high across lipid species (97.6% ± 2.0%), whereas in the mother the lipids were substantially less deuterated (60.6% ± 26.4%) with levels varying among lipid species and subspecies. The mass distribution pattern of deuterium-containing isotopologues indicated the fraction (in %) of each lipid (sub-)species having one or more deuteriums incorporated: in the D-pups, the pattern was always bell-shaped, and the average number of D atoms ranged from a low of ∼4 in fatty acid to a high of ∼9 in cerebroside. By contrast, in D-dam most lipids had more complex, overlapping distributions that were weighted toward a lower average number of deuteriums, which ranged from a low of ∼3–4 in fatty acid and in one species of sulfatide to a high of 6–7 in cerebroside and sphingomyelin. The consistently high level of deuteration in D-pups can be attributed to their *de novo* lipogenesis during gestation and rapid, postnatal myelination. The widely varying levels of deuteration in D-dam, by contrast, likely depends on the relative metabolic stability of the particular lipid species during myelin maintenance. Our current findings demonstrate that stably-incorporated D label can be detected and localized using neutron diffraction in a complex tissue such as myelin; and moreover, that MS can be used to screen a broad range of deuterated lipid species to monitor differential rates of lipid turnover. In addition to helping to develop a comprehensive understanding of the *de novo* synthesis and turnover of specific lipids in normal and abnormal myelin, our results also suggest application to studies on myelin proteins (which constitute only 20–30% by dry mass of the myelin, vs. 70–80% for lipid), as well as more broadly to the molecular constituents of other biological tissues.

## Introduction

Diffraction studies of myelin membranes and their molecular constituents have helped to characterize the molecular organization that underlies myelin stability and instability ([1] and references therein). These issues are important because of myelin’s essential role in fast, efficient nerve conduction and involvement in a variety of neurological conditions caused by or resulting in demyelination [2]. Because myelin is a multilamellar stack of membranes, lamellar diffraction has been especially useful in providing information about the general distribution of lipids, protein, and water across the membranes [3] and also about the localization of specific lipids (e.g., cholesterol [4]; phosphatidylethanolamine plasmalogen [5]) and proteins (e.g., myelin basic protein and P0 glycoprotein [6]). The majority of diffraction studies of myelin use X-rays; however, neutron diffraction offers a particular advantage owing to the large difference in scattering lengths between hydrogen (H) and deuterium (^2^H or D), and hence between H_2_O and D_2_O (deuterium oxide; heavy water). That deuterium can be localized by neutron diffraction suggests the usefulness of both deuterium exchange and deuterium labeling. With H→D exchange, simple diffusion can result in a distinct, detectable change in the relative amounts of H_2_O and D_2_O in the aqueous compartments of multilamellar myelin, such as membrane regions that have accessible, exchangeable hydrogens. Thus, the distribution of water across the membrane bilayer as well as the kinetics of H_2_O/D_2_O exchange can be directly determined using these isotopologues of water [7,8] that cannot be distinguished using X-rays. Neutron diffraction studies using H-D exchange have confirmed the thickness of the water-excluding hydrocarbon domain deduced from X-ray diffraction [4,9], and the rate of D-H exchange in the water-accessible spaces [10]. Deuterium labeling that is non-covalent can be carried out by diffusing in a deuterated compound to localize that moiety, e.g., deuterated DMSO [11]; whereas deuterium labeling via covalent bonding is undertaken by the introduction of deuterated myelin constituents (e.g., D-cholesterol [12]), or precursors of the constituents (D-choline, D.L. Worcester, personal communication, 2019), or by providing D_2_O-spiked drinking water to laboratory animals [13,14]. In the case of highly-specific metabolic precursors, the labeled compound must first pass through either the blood-nerve barrier (for PNS myelin) or the blood-brain barrier (for CNS myelin) before it can target a particular molecule. By contrast, introducing deuterium via D_2_O-spiked drinking water ensures that this non-specific label is incorporated metabolically into the full complement of the body’s molecular constituents that comprise its cells and tissues.

In the current study, we administered D_2_O to a pregnant mouse via drinking water, and continued this protocol two months postpartum for both the dam and her pups. Neutron diffraction from sciatic nerves (reported here) and spinal cords (not shown) between H_2_O and D_2_O-fed dams (H-dam, D-dam) and their pups (H-pup, D-pup) revealed significant differences in the diffraction patterns. Neutron scattering length density profiles of the myelin membranes showed a prominent increase in scattering density in the middle of the membrane bilayer, consistent with incorporation of deuterium into the lipid hydrocarbon. This novel localization by neutron diffraction of deuterium that was metabolically-incorporated into lipid was subsequently confirmed by mass spectrometry (MS) analysis of trigeminal nerve lipids, which moreover identified the lipid species as phosphatidylethanolamine (PE), phosphatidylcholine (PC), phosphatidylserine (PS), cerebroside (HexCer), sulfatide (SHexCer), sphingomyelin (SM), phosphatidylinositol (PI), triacylglycerol (TG), diglycerol (DG), and fatty acid (FA).

Whereas the % of lipids with at least one deuterium averaged ∼98 ± 2% for D-pups, the % of lipids for D-dam indicated variable deuteration, averaging ∼61 ± 26%, and moreover, depended on lipid species and subspecies. The consistent, uniformly high incorporation of D into lipids for the D-pups vs. that for D-dam can be explained by the *de novo* biosynthesis and myelin assembly in the former vs. differential turnover of lipids in mature myelin [13]. In summary, we demonstrated that neutron diffraction together with mass spectrometry can be used to detect, localize, and identify deuterium that is stably-incorporated via normal metabolism into the rapidly- and newly-synthesized myelin as well as into myelin undergoing metabolic maintenance where lipid turnover is more targeted.

## Materials and methods

### Reagents and standards

Methyl-tert-butyl-ether (MTBE) and chloroform were purchased from Sigma Aldrich; methanol was from Fisher Science Education; ammonium acetate was from Fluka; and D_2_O (99.9%) was from Sigma-Aldrich Corp and from Eurisotop.

### Specimens

All animal procedures were conducted at the Boston College Animal Care Facility in accordance with protocols approved by the Institutional Animal Care and Use Committee at Boston College. Three timed-pregnant C57BL/6 mice were obtained from Charles River Laboratories (Wilmington, MA), housed on a 12-hour light/dark cycle, and provided *ad libitum* access to food and water. At gestational day 12, when the dams were ∼10 weeks old, the mice were separated into treatment groups, in which their drinking water was replaced with water containing 15% (D-mice) or 0% (control, H-mice) D_2_O. Mice were maintained as such through birth and until postnatal day 7, at which point the volume fraction of D_2_O in their drinking water was increased to 20% (D-mice). At postnatal day 13, their drinking water was again increased to 30% volume fraction of D_2_O. Pups were weaned at postnatal day 28, and all mice were maintained on 30% D_2_O (D-mice) or 0% D_2_O (control, H-mice) until the pups reached postnatal day 55, at which point all mice (including the dams at 131 do) were sacrificed (see S1 Fig) and tissues were harvested as described in the following. The chosen level of D_2_O in drinking water was below the threshold for toxicity for mice [13,15,16].

### Sample preparation

Mice were sacrificed by CO_2_ asphyxiation, followed by decapitation. Brains (including optic nerves) and trigeminal nerves (which, like sciatic nerves, are peripheral nerves) were removed, flash frozen, and stored at -80°C until further prepared for MS. Sciatic nerves and sagittally-bisected segments of spinal cord were removed for analysis by neutron diffraction. Sciatic nerves were tied off at both ends with silk suture and placed under gentle tension on plastic sample holders, while spinal cord segments were laid flat, adhering naturally to filter paper. These samples were immersion-fixed in 1% paraformaldehyde and 1.5% glutaraldehyde in 0.12 M phosphate buffer pH 7.4 overnight at room temperature (RT). Fixed samples were then equilibrated against phosphate-buffered saline (PBS; 5 mM phosphate buffer, 154 mM NaCl, pH/pD 7.4) of varying D_2_O content (0–100%) overnight and loaded into thin-walled (20 μm) quartz capillary tubes that were 0.7 mm-diameter for sciatic nerves and 1.5 mm-diameter for spinal cords (Charles Supper Company; Natick, MA) containing PBS with the same D_2_O fraction. The capillaries were sealed with wax and enamel and stored at RT until analyzed by neutron diffraction (S1 Table). During the diffraction experiments, some samples were examined under more than one set of conditions by removing the tissue from the capillary tube, equilibrating it overnight against PBS containing a different volume fraction of D_2_O, and then resealing it in a fresh capillary tube.

### Neutron diffraction

The experiments were carried out on the D16 instrument at the Institut Laue-Langevin (ILL; Grenoble, France). The monochromatic neutron beam (λ = 4.55 Å, Δλ/λ = 0.01) was vertically focused by nine Highly Oriented Pyrolytic Graphite (HOPG) crystals (average mosaic 0.4°), and to optimize sample illumination and angular resolution two pairs of collimating slits were used. The resulting beam size at the sample was 1 mm (horizontal) x 10 mm (vertical). The monochromator-to-sample distance was 2980 mm and the sample-to-detector distance was 955 mm. Diffraction patterns were collected using the Millimeter Resolution Large Area Neutron Detector (MiLAND), a flat, high pressure ^3^He neutron detector with an area of 320 mm x 320 mm and a “pixel” resolution of 1 mm x 1 mm. For all experiments, the detector angle relative to the incident beam (Gamma axis) was set to 5.2°, so that it could collect diffraction patterns in a 2θ-range of -4° to 14.6°, corresponding to a range in *q* from -0.096 Å^-1^ to 0.35 Å^-1^, which included the attenuated direct beam, each of the 1^st^ and 2^nd^ order reflections on both sides of the direct beam, and up to the 7^th^ order reflection on one side of the beam.

All neutron diffraction experiments were performed at RT and pressure, consistent with our previous diffraction experiments at the ILL [8,10]. The entubed tissues were mounted and aligned in a dedicated sample changer; and exposure times for data collection ranged from 1–3 h depending on water contrast and whether the sample was sciatic nerve or spinal cord. In the current paper, we report on our structural analysis of the sciatic nerves.

Data refinement was performed using the ILL in-house software LAMP ([17]; http://www.ill.eu/instruments-support/computing-for-science/data-analysis). First, two-dimensional diffraction patterns were normalized to the incident neutron beam flux to account for variations in beam intensity and exposure time between samples. Second, they were normalized to a detector calibration file containing the flat incoherent signal from water, which is used to correct the observed intensities for pixel efficiency and solid angle. Third, they were corrected for the attenuation of the direct beam by the sample (transmission). Finally, background patterns from empty quartz capillaries and instrument background were collected and subtracted from each sample pattern, and normalization to the sample thickness was applied. The corrected two-dimensional patterns were then integrated vertically (along the nerve axis) to produce one-dimensional diffraction patterns (Fig 1).

**Fig 1.**
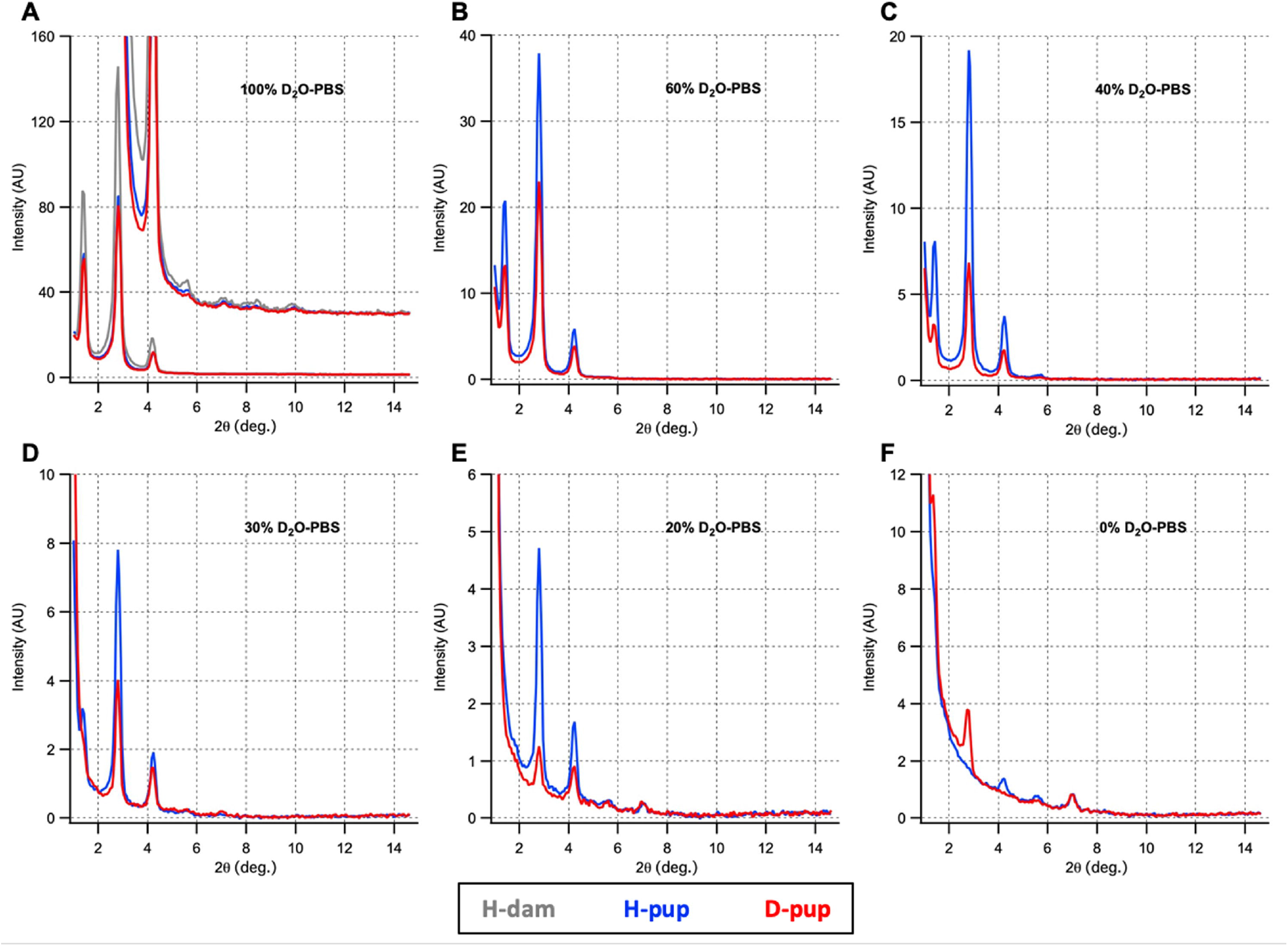
Neutron diffraction spectra from the (fixed) sciatic nerves of 2-month old pups that had gestated and then been raised on either normal drinking water or on a mixture of D_2_O in water (see Materials and methods for protocol). Dissected nerves were equilibrated in 0%–100% D_2_O-PBS. The scattering intensity, in arbitrary units (AU), is plotted as a function of scattering angle 2□ in degrees. At 100%–30% D_2_O-PBS (A-D), Bragg orders 1–3 are clearly evident at 2□ ≈ 1.4, 2.8, and 4.2, respectively. (Some of the higher Bragg orders (4–7) are seen here in panels A,C,E,F at 2□ ≈ 5.6, 7.0, 8.4, and 9.8.) For each type of pup, the relative intensities change systematically as the %D_2_O changes, and these changes differ between the D-pups (red) vs. H-pups (blue). (A) includes the diffraction pattern from H-dam (gray), the non-deuterated mother of the H-pups, showing the similarity in relative intensities of the Bragg peaks between mother and pup, but nearly twice as strong for the mother. (B-F) show that as the amount of diffusible D_2_O decreases, the diffracted intensities change more dramatically in the deuterated pup (D-pup) compared to the non-deuterated pup (H-pup), owing to localization of the non-diffusible and stable, metabolically-incorporated deuterium in the former. Diffraction from H-dam in 100%, 40%, and 20% D_2_O-PBS is shown in S4A Fig.

Peak positions, peak widths, and integrated intensities above background were determined using MagicPlot software (Magicplot Systems, LLC; https://magicplot.com/). Then, for all samples the myelin period (*d*) was calculated according to:

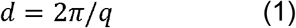

where *q* is the scattering vector given by:

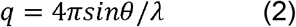

where λ is the neutron wavelength, and θ is one-half the scattering angle. For a precise determination of the myelin spacing, we plotted the *q*_*h*_ peak positions of each sample vs. order *h*. Combining Eq. (2) with Bragg’s law (*hλ* = 2*dsinθ*) yielded the linear dependency between *q*_*h*_ and *h*; and a linear fit to the *q*_*h*_ *vs. h* data provided the slope and, therefore, the *d*-spacing (S2 Fig).

Structure amplitudes (F(*h*)) were calculated from the peak intensities (*I(h)*) using the equation

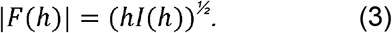

Linear regression analysis was performed between *F(h)* and %D_2_O. Phases were assigned to structure factors according to [7,8,18]. Average neutron scattering density profiles were calculated using *F(h)* according to

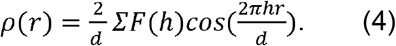

Owing to changes in the widths of the water-accessible spaces in fixed myelin [18], determining the phases for the structure amplitudes measured for the fixed myelin analyzed here involved revising the strategy used previously for native (unfixed) myelin [7,11]. Briefly, for tissues from H-dam and H-pups that were not exposed to deuterium-spiked water, a step-function approximation of the contrast between the hydrocarbon of the membrane bilayer and the non-hydrocarbon layers was used to represent the scattering density profile of the myelin: for myelin in 100% D_2_O-PBS, the major contrast is between the hydrocarbon and the water layers at the cytoplasmic and extracellular appositions of the membrane multilayers; and for myelin in 0% D_2_O-PBS, the major contrast is between the hydrocarbon and the lipid polar groups. For the step-function approximation, we measured the widths of these layers from the X-ray scattering density profile of glutaraldehyde-fixed myelin of mouse sciatic nerve, in which the membrane bilayer dimensions have remained constant, whereas the widths of the cytoplasmic and extracellular separations have decreased and increased, respectively [18]. We revised the step-function model for unfixed myelin [7,11] by incorporating these altered dimensions of the aqueous spaces (S3 Fig). The modified step-function model was used to calculate membrane pair transforms for the fixed myelin in 100%-D_2_O-PBS and in 0% D_2_O-PBS (Fig 2, continuous curves). The agreement between the observed scattering amplitudes and those predicted by the transform sampled at *h*/186.1 Å (for *h*=1–5) established the phases for the neutron diffraction data (Fig 2, data points; Table 1). Moreover, linear regression between *F(h)* and %D_2_O (Fig 3) supported the phase assignments based on the step-function models.

**Table 1.**
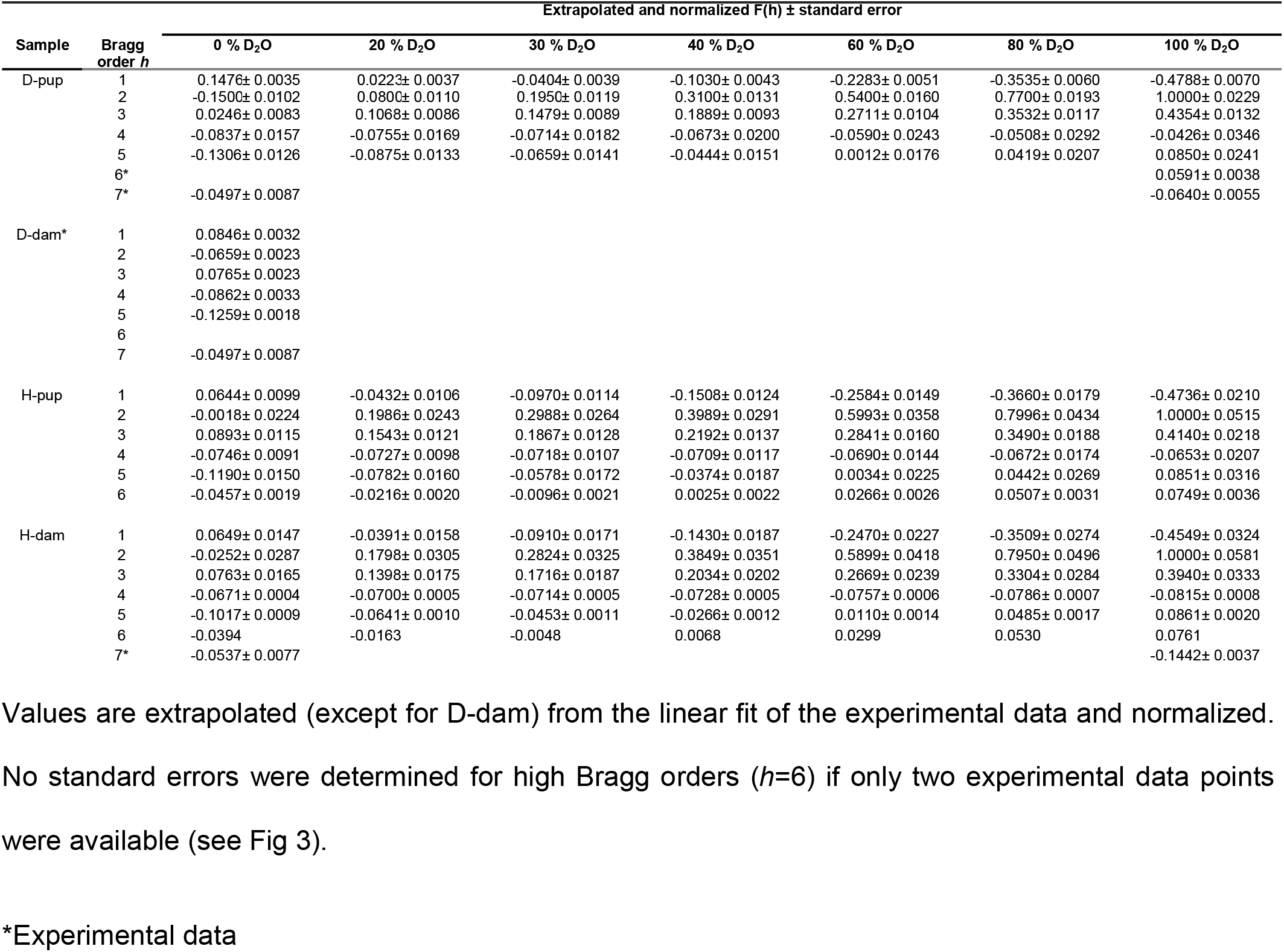
Structure-factor for D-pup, D-dam, H-pup and H-dam vs. %D_2_O.

| Sample | Bragg order $h$ | Extrapolated and normalized $F(h) \pm$ standard error | | | | | | |
| --- | --- | --- | --- | --- | --- | --- | --- | --- |
|  |  | 0 % D<sub>2</sub>O | 20 % D<sub>2</sub>O | 30 % D<sub>2</sub>O | 40 % D<sub>2</sub>O | 60 % D<sub>2</sub>O | 80 % D<sub>2</sub>O | 100 % D<sub>2</sub>O |
| D-pup | 1 | 0.1476± 0.0035 | 0.0223± 0.0037 | -0.0404± 0.0039 | -0.1030± 0.0043 | -0.2283± 0.0051 | -0.3535± 0.0060 | -0.4788± 0.0070 |
|  | 2 | -0.1500± 0.0102 | 0.0800± 0.0110 | 0.1950± 0.0119 | 0.3100± 0.0131 | 0.5400± 0.0160 | 0.7700± 0.0193 | 1.0000± 0.0229 |
|  | 3 | 0.0246± 0.0083 | 0.1068± 0.0086 | 0.1479± 0.0089 | 0.1889± 0.0093 | 0.2711± 0.0104 | 0.3532± 0.0117 | 0.4354± 0.0132 |
|  | 4 | -0.0837± 0.0157 | -0.0755± 0.0169 | -0.0714± 0.0182 | -0.0673± 0.0200 | -0.0590± 0.0243 | -0.0508± 0.0292 | -0.0426± 0.0346 |
|  | 5 | -0.1306± 0.0126 | -0.0875± 0.0133 | -0.0659± 0.0141 | -0.0444± 0.0151 | 0.0012± 0.0176 | 0.0419± 0.0207 | 0.0850± 0.0241 |
|  | 6* |  |  |  |  |  |  | 0.0591± 0.0038 |
|  | 7* | -0.0497± 0.0087 |  |  |  |  |  | -0.0640± 0.0055 |
| D-dam* | 1 | 0.0846± 0.0032 |  |  |  |  |  |  |
|  | 2 | -0.0659± 0.0023 |  |  |  |  |  |  |
|  | 3 | 0.0765± 0.0023 |  |  |  |  |  |  |
|  | 4 | -0.0862± 0.0033 |  |  |  |  |  |  |
|  | 5 | -0.1259± 0.0018 |  |  |  |  |  |  |
|  | 6 |  |  |  |  |  |  |  |
|  | 7 | -0.0497± 0.0087 |  |  |  |  |  |  |
| H-pup | 1 | 0.0644± 0.0099 | -0.0432± 0.0106 | -0.0970± 0.0114 | -0.1508± 0.0124 | -0.2584± 0.0149 | -0.3660± 0.0179 | -0.4736± 0.0210 |
|  | 2 | -0.0018± 0.0224 | 0.1986± 0.0243 | 0.2988± 0.0264 | 0.3989± 0.0291 | 0.5993± 0.0358 | 0.7996± 0.0434 | 1.0000± 0.0515 |
|  | 3 | 0.0893± 0.0115 | 0.1543± 0.0121 | 0.1867± 0.0128 | 0.2192± 0.0137 | 0.2841± 0.0160 | 0.3490± 0.0188 | 0.4140± 0.0218 |
|  | 4 | -0.0746± 0.0091 | -0.0727± 0.0098 | -0.0718± 0.0107 | -0.0709± 0.0117 | -0.0690± 0.0144 | -0.0672± 0.0174 | -0.0653± 0.0207 |
|  | 5 | -0.1190± 0.0150 | -0.0782± 0.0160 | -0.0578± 0.0172 | -0.0374± 0.0187 | 0.0034± 0.0225 | 0.0442± 0.0269 | 0.0851± 0.0316 |
|  | 6 | -0.0457± 0.0019 | -0.0216± 0.0020 | -0.0096± 0.0021 | 0.0025± 0.0022 | 0.0266± 0.0026 | 0.0507± 0.0031 | 0.0749± 0.0036 |
| H-dam | 1 | 0.0649± 0.0147 | -0.0391± 0.0158 | -0.0910± 0.0171 | -0.1430± 0.0187 | -0.2470± 0.0227 | -0.3509± 0.0274 | -0.4549± 0.0324 |
|  | 2 | -0.0252± 0.0287 | 0.1798± 0.0305 | 0.2824± 0.0325 | 0.3849± 0.0351 | 0.5899± 0.0418 | 0.7950± 0.0496 | 1.0000± 0.0581 |
|  | 3 | 0.0763± 0.0165 | 0.1398± 0.0175 | 0.1716± 0.0187 | 0.2034± 0.0202 | 0.2669± 0.0239 | 0.3304± 0.0284 | 0.3940± 0.0333 |
|  | 4 | -0.0671± 0.0004 | -0.0700± 0.0005 | -0.0714± 0.0005 | -0.0728± 0.0005 | -0.0757± 0.0006 | -0.0786± 0.0007 | -0.0815± 0.0008 |
|  | 5 | -0.1017± 0.0009 | -0.0641± 0.0010 | -0.0453± 0.0011 | -0.0266± 0.0012 | 0.0110± 0.0014 | 0.0485± 0.0017 | 0.0861± 0.0020 |
|  | 6 | -0.0394 | -0.0163 | -0.0048 | 0.0068 | 0.0299 | 0.0530 | 0.0761 |
|  | 7* | -0.0537± 0.0077 |  |  |  |  |  | -0.1442± 0.0037 |

\*Experimental data

**Fig 2.**
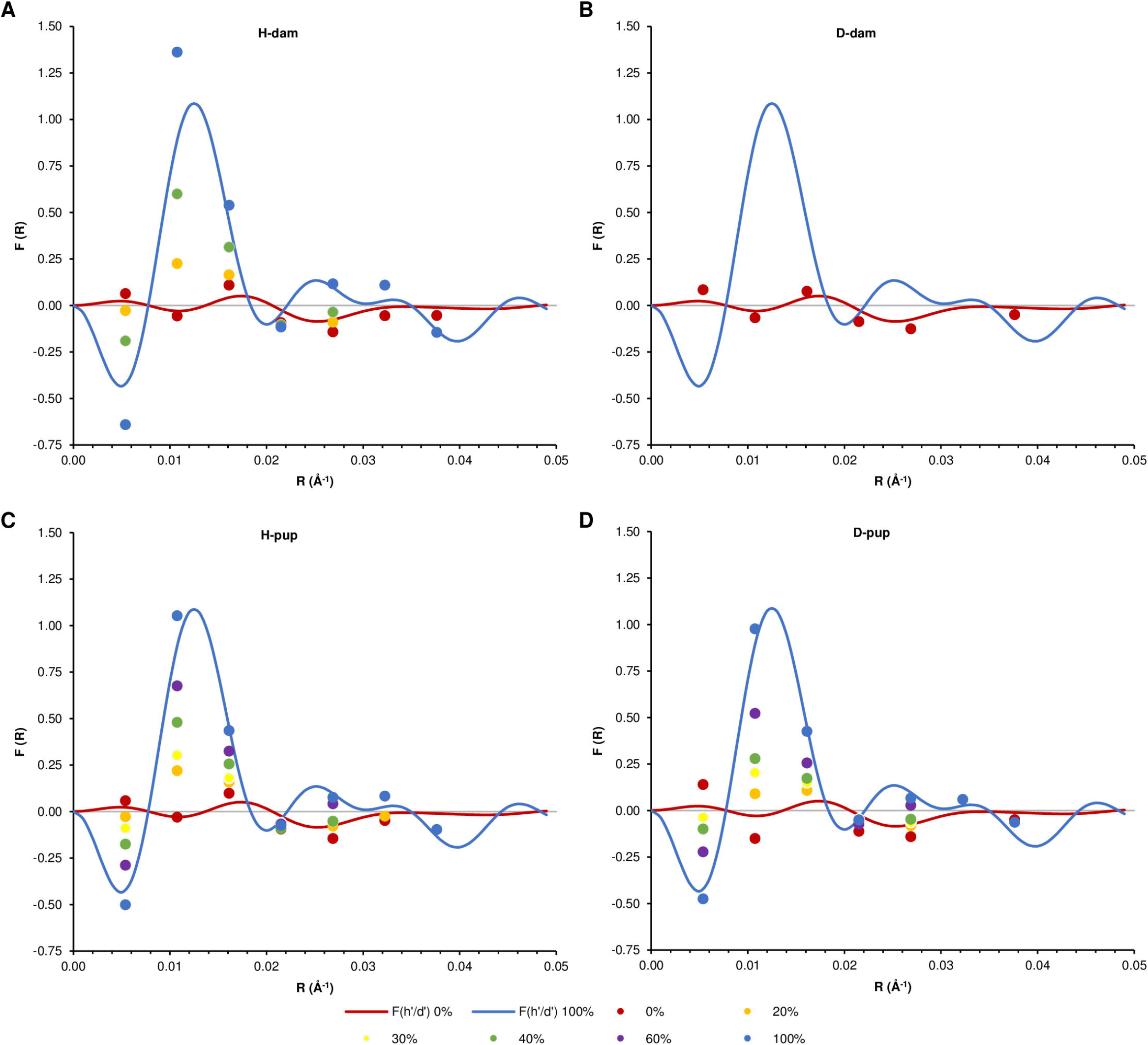
Determination of phases from step-function. The continuous transforms *F(R)* for H-dam at 100% D_2_O-PBS and at 0% D_2_O-PBS were calculated as described (see Materials and methods: Neutron diffraction, and S3 Fig) and used to predict phases for the structure amplitudes calculated from the Bragg orders. Experimental data points for H-dam (A), D-dam (B), H-pup (C), and D-pup (D) at different amounts of D_2_O in PBS (0%, 20%, 30%, 40%, 60%, 80%, and 100%) were mapped onto the continuous transform, and showed a systematic change in *F(h)* (Fig 3).

**Fig 3.**
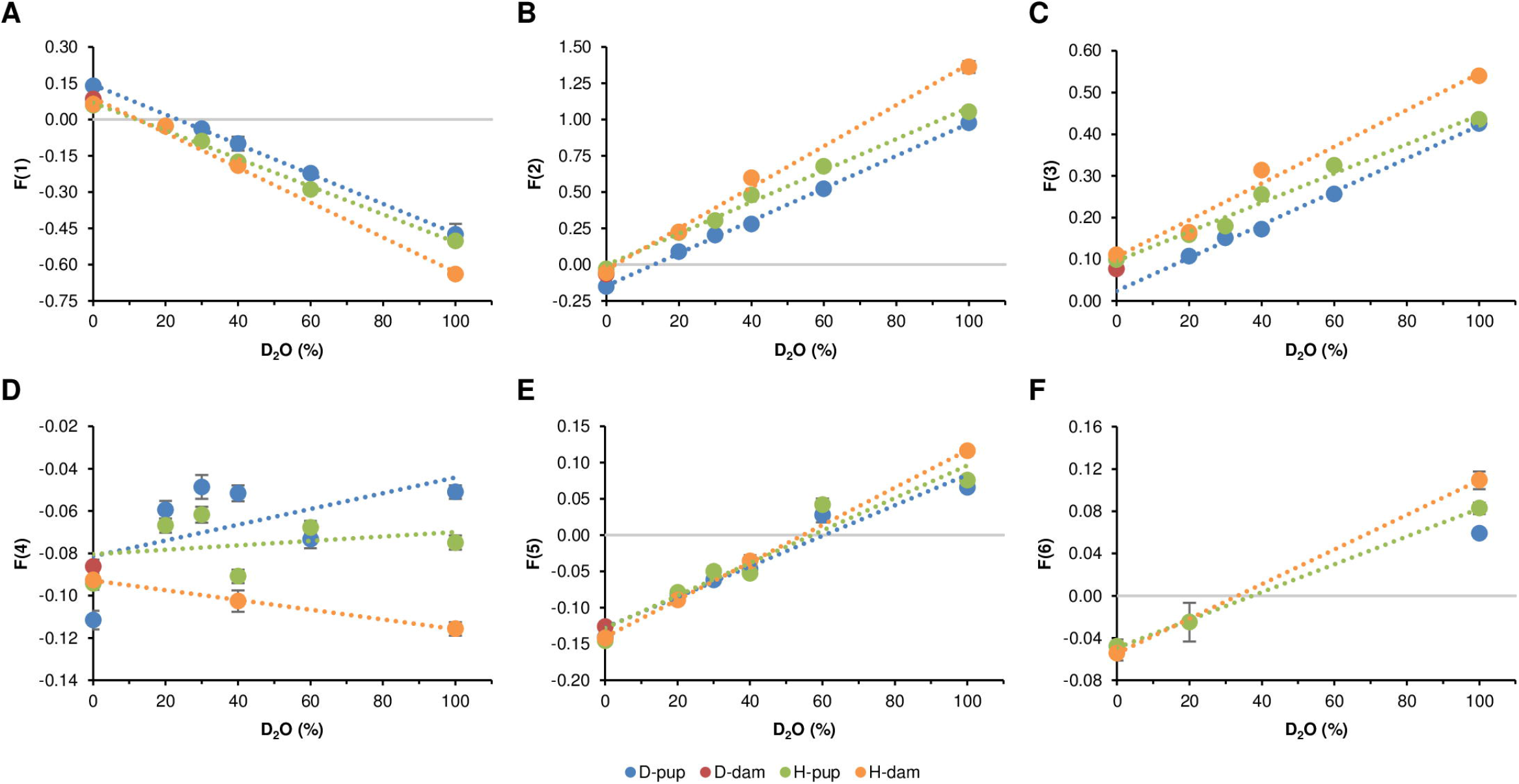
Structure-factor amplitudes with assigned phases vs. %D_2_O. Symbols represent experimental replicates and lines the linear dependence of *F(h)* on %D_2_O. □*F* is calculated by using □*F* = *F* * (0.5 * □*I/I*), where □*I* is the standard deviation in fitting backgrounds and intensities of the Bragg orders.

## Mass spectrometry

### Homogenization and lipid extraction

Trigeminal nerves were homogenized in ice-cold 0.1% ammonium acetate in a tissue grinder. Methanol (375 µl) was added to 50 µl homogenate, which was placed into a glass tube with a screw cap, and the tube was vortexed. Further, 1.25 ml MTBE was added and the mixture was incubated for 1 h at RT under agitation. Phase separation was induced by adding 312.5 µl milliQ water. Upon 10 min incubation at RT, the sample was centrifuged at 1000 g for 10 min. The upper organic phase was collected and dried in a vacuum centrifuge at RT. Dried samples were stored at -20°C until further use.

### Thin-layer chromatography

High-performance thin-layer chromatography (HPTLC) was used to analyze polar and neutral lipids as described previously [19]. Lipids were spotted on Partisil LHPK Silica gel 60A plates (Whatman): 300□µg nerve dry weight for polar lipids, and 300□µg nerve dry weight for neutral lipids. A lipid standard mix (polar lipids: non-hydroxy cerebroside, hydroxy cerebroside, non-hydroxy sulfatide, hydroxy sulfatide, phosphatidylethanolamine, phosphatidylinositol, phosphatidylserine, phosphatidylcholine, sphingomyelin; neutral lipids: cholesterol esters, triglycerides, diglycerides, cholesterol, free fatty acids, monoglyceride) was spotted on plates at 2.4 µg. Polar lipids were separated with methyl acetate:1-propanol:chloroform:methanol:0.25% KCl:acetic acid (25:25:25:10:9:0.3 by volume). Neutral lipid were first separated with chloroform:methanol:acetic acid (93.1:1.9:0.1 by volume) to a height of 8 cm and then separated to the top with hexane:ethyl ether:acetic acid (89.3:5.7:0.1 by volume). Polar and neutral lipids were visualized with 10% CuSO_4_, 8% H_3_PO_4_ and heating at 160°C for 13 min. The distribution and relative intensity of the charred lipids (S10 Fig) closely resembled our previous data from mouse nerve lipids [19].

### Lipid analysis by mass spectrometry

Lipid extracts were resuspended in 1 ml methanol/chloroform (1:1, v/v) and an aliquot of 150 µl was re-dried in a vacuum centrifuge. Finally, the dried lipid sample was reconstituted with 200 µl mobile phase A (acetonitrile: water, 60:40, v/v) and mobile phase B (isopropanol/ acetonitrile, 90:10, v/v) using a ratio of 60:40. 100 µl was transferred into an MS vial. Samples were separated using a CORTECS™ C_18_ column (Waters, 90 Å, 2.7 µm, 2.1 mm X 100 mm). Lipid extracts were analyzed with an Agilent 6220 mass spectrometer coupled with an Agilent 1200 series HPLC system in both positive and negative ion modes. Instrument calibration was accomplished using a series of Hexakis compounds (Agilent Technologies, Santa Clara, CA: ESI-L Low Concentration Tuning Mix, Part No. G1969-85000, Lot No. LC21022V); and the reference solution (Agilent: API-TOF Reference Mass Solution Kit, Part No. G1969-85001, Lot No. LB48019) consisted of purine (C_5_H_4_N_4_; CAS#120-73-0 and hexakis (Hexakis(1h,1h,3h-tetrafluoropropoxy)phosphazine (C_18_H_18_F_24_N_3_O_6_P_3_; CAS#58943-98-9). 10 µl of each experimental sample was injected on the column at a flow rate of 350 µl/min, 65°C column temperature, and with the following gradient: 0 min 40% mobile phase B (see above), 1 min 40% B, 11 min 100% B, 15 min 100% B, 16 min 40% B, with 4 min equilibration. Mass spectrometry parameters were as follows: *m/z* range 100–3000, fragmentor 350 V, skimmer 65 V, gas temp 325°C, drying gas 8 l/min, nebulizer 45 psig, capillary 4000 V. As an example, S11 Fig shows the chromatogram for a lipid extract (upper panel) with the mass spectra determined for the same retention time range for the control (H-dam) and treated (D-dam) (middle and lower panels, respectively).

Identification of lipid monoisotopes in the MS spectra of samples from the H-dam and H-pups was initially based on *m/z* matching using MS-LAMP [20]; and the identification of *m/z* was vetted by comparison with the updated, curated data in two databases (The LIPID MAPS® Lipidomics Gateway, https://www.lipidmaps.org/ [21,22] and SwissLipids, https://www.swisslipids.org/#/advanced [23]) (Table 2). We further vetted the matching of our experimental *m/z* values to those of lipid species reported specifically for mouse neural tissues (sciatic nerve [24]; cervical dorsal root ganglia [25]; CNS myelin [26]; optic nerve [27]; brain tissues [28–30]; cerebellum [31]). We also used the extensive tabulated data for rat (adult, male Wistar) sciatic nerve lipids that were analyzed by imaging MS [32]. In our current study, the conditions we used for analyzing the lipids were not sufficient to detect cholesterol, a major myelin constituent, which is usually cationized via alkaline hydrolysis and detected, for example, as ammoniated, lithiated, or sodiated adducts [33]. Spectral accuracy of our original data was determined using MassWorks (Version 6,0,0,0) software from Cerno Bioscience (LasVegas, NV 89144) [34,35]. (The MassWorks software also corrected automatically for the presence in the deuterated lipids of the natural isotope abundance that was detected in the non-deuterated samples (H-dam and H-pup spectra, see below). Subsequently, we used MassWorks to identify the corresponding deuterated isotopologues in the isolated lipids from the D-mice based on the vetted undeuterated monoisotopes for the isolated lipids from the H-mice, and to quantitate the relative abundance (%) Ai of each isotopologue. The average level of deuteration was calculated by summing these relative percentages (i.e., %D = Σ_i>0_ A_i_). From the isotopologue distribution we also calculated the weighted average number (and standard error) of deuteriums for each molecular species of lipid [36]. The stated uncertainties in the quantitation of the data are standard deviations, unless indicated otherwise.

**Table 2.**
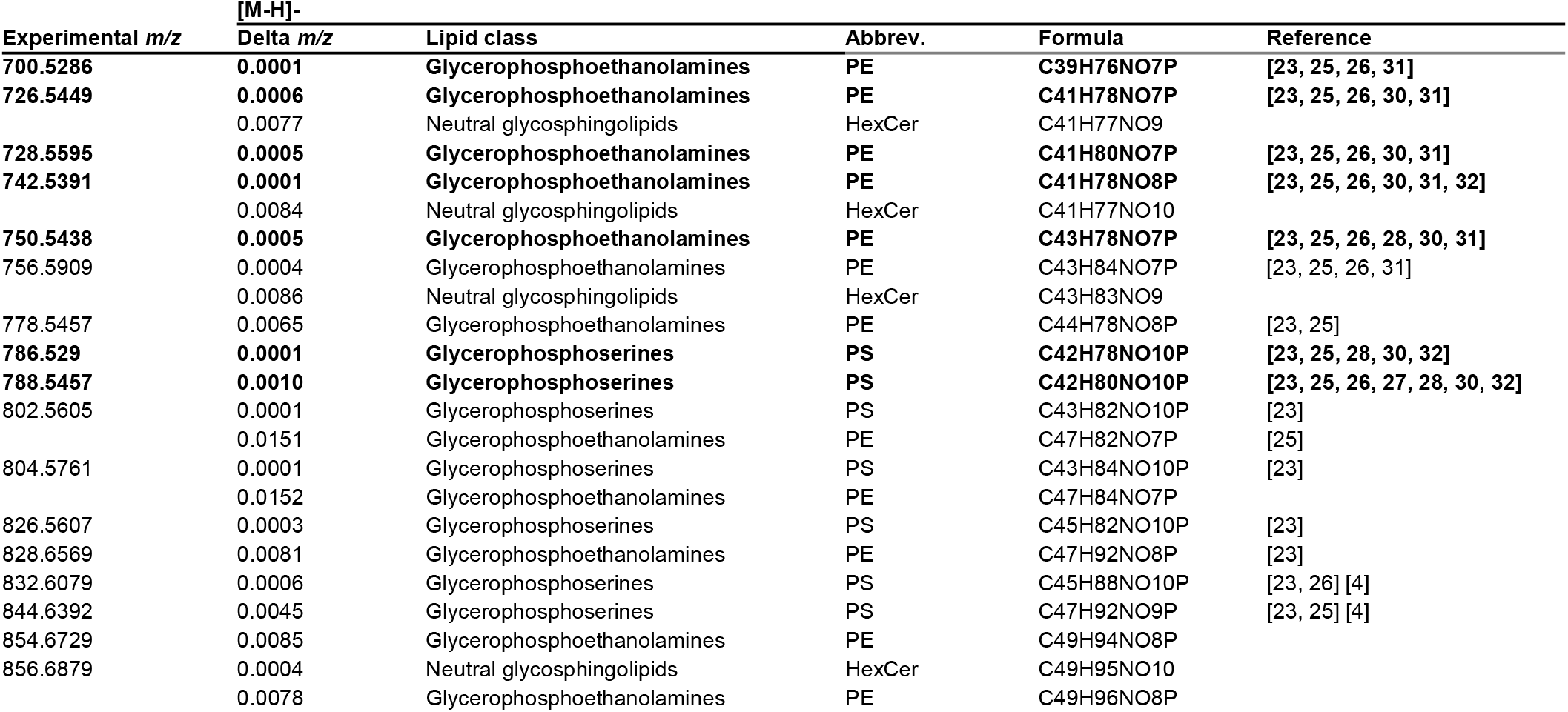
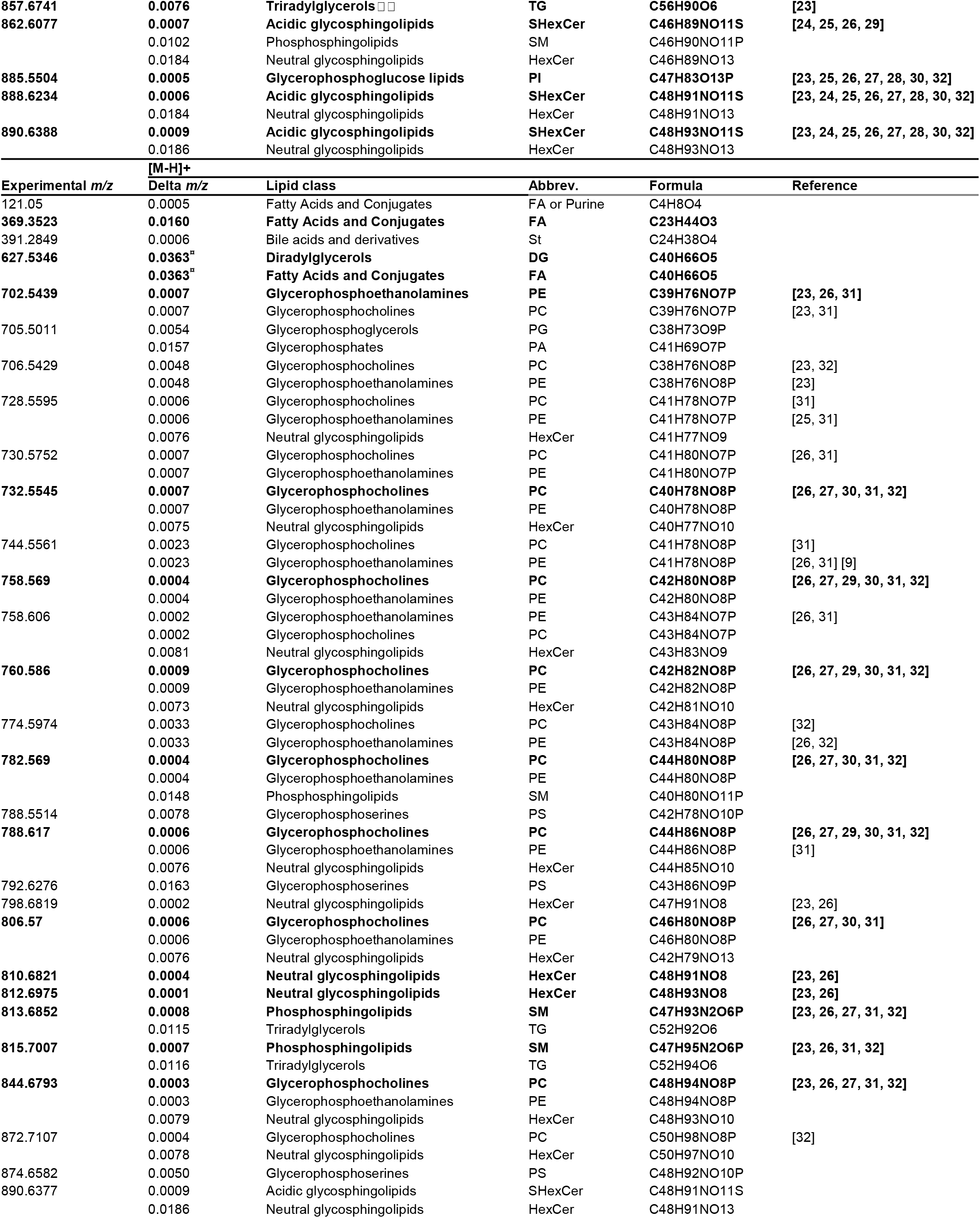
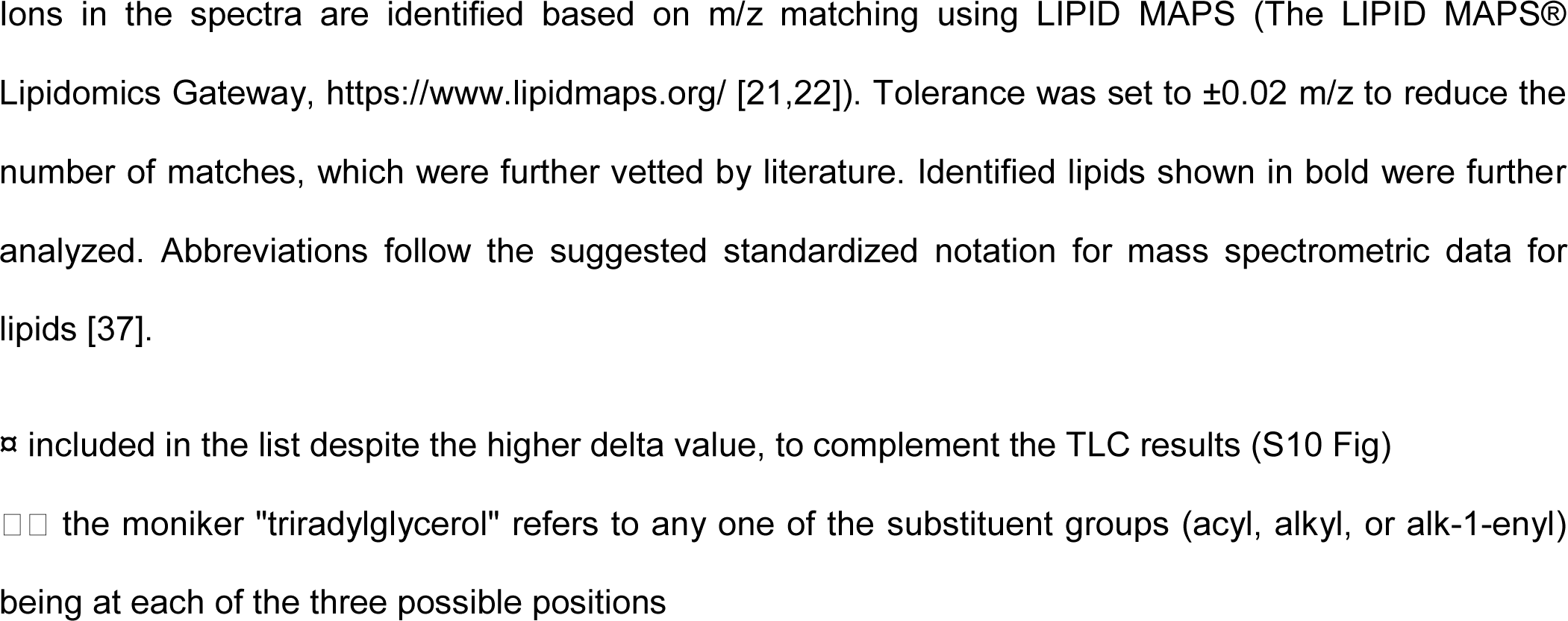
Identified lipids in mouse trigeminal nerves using LC-MS.

## Results and discussion

### Use of fixed tissue for diffraction

Owing to the logistical complexity in raising pregnant dams and their offspring on a strict regimen of defined D_2_O-H_2_O drinking water over a period of several months at the ESRF’s animal facility that was accessible to us at the ILL (Grenoble), we undertook the long-duration, preparative portion of the experiment at Boston College. As a result, to maintain as well as possible the structural integrity of the tissues during their transit to Grenoble from Boston, we fixed the tissue in aldehyde solutions and stored them in PBS (as described above, Materials and methods: Sample preparation). Previous experiments that used X-ray diffraction to document the structural changes in myelin during processing for electron microscopy show that the initial fixation step results in relatively small changes in myelin periodicity, membrane packing, and electron density distribution compared to the subsequent steps in tissue processing [18,38]. Whereas the diffracted intensity is nearly as strong as from unfixed nerves, there are, however, changes in the positions of the Bragg orders and their relative intensities that arise from the narrowing of the cytoplasmic apposition and widening of the extracellular space between membranes. In spite of such well-defined changes in membrane packing caused by chemical fixation, numerous X-ray diffraction studies indicate that intrinsic differences between different types of myelin that are identically fixed may still be detected [38–43].

The neutron diffraction patterns for unfixed (data from [8]) and fixed (H-dam, this study) mouse sciatic nerves in 100% D_2_O-PBS showed a shift in the positions of the Bragg orders, which is accounted for by the increase in myelin periodicity to 186.1 ± 0.8 Å (n=17) from 175.3 ± 2.2 Å (n=5; [8]). The first three Bragg orders, which have the strongest intensities at 100% D_2_O in both unfixed (see Fig 1c in [8]) and fixed myelin (this manuscript, Fig 1A), dominate the patterns (S4B Fig). The main difference is the stronger 1^st^ order intensity in fixed compared to unfixed myelin. Among the higher orders, the main difference is the increase in the 5^th^ and decrease in the 6^th^ order in the fixed compared to the unfixed. These differences in periodicities and intensities between unfixed and fixed myelin are mostly accounted for by the change in packing of the membranes in the myelin array—namely, the widening of the extracellular space and narrowing of the cytoplasmic space between membrane pairs (S5 Fig). By contrast, the myelin that had been deuterated via metabolic incorporation from D_2_O (and then fixed) showed indistinguishable periodicities (S2 Fig) but different intensities compared to the (fixed) non-deuterated tissue (Fig 1). This suggests that there was an appreciable and specific localization of deuterium that had been incorporated stably into the membrane structure.

### Systematic changes in scattering density of membrane profiles consistent with deuteration of lipids

To reveal the localization of deuterium in the myelin membrane arrays, we calculated the scattering length density profiles from the phases assigned to the structure amplitudes (see Materials and methods: Neutron diffraction). This determination involved use of the step-function defined by the water-accessible spaces of myelin (S3 Fig), calculation of membrane transforms from the models to predict structure amplitudes (Fig 2), and linear regression between the structure amplitudes and %D_2_O in the water spaces (Fig 3) to confirm phase assignments.

The profiles showed that the H-pups and the H-dam corresponded closely in scattering density when the nerves were equilibrated in 0% D_2_O-PBS or in 100% D_2_O-PBS (Fig 4A; see also S8 Fig): i.e., the centers of the membrane bilayers at ∼40 □, the neighboring peaks at ∼20 □and 60 □, and the inter-bilayer, water-accessible spaces centered at 0 □and ∼90 □superimposed in both position (x-axis) and level (y-axis). By comparison, after equilibration of nerves in 0% D_2_O-PBS the scattering density profiles for the D-pups and D-dam superimposed at the peak and trough positions (x-axis), but the density levels (y-axis) for D-pup were significantly higher than those of D-dam except in the water-accessible spaces centered at 0 □and ∼90 □between membrane bilayers (Fig 4B). Thus, myelin from young mice that had been exposed to deuterium via D_2_O during gestation showed a more pronounced increase in scattering density in the membrane bilayer region compared to their mother, who had also been exposed to the same D_2_O-spiked drinking water. The greater incorporation of deuterium in the pups is consistent with the greater extent of myelination during gestation and development than during the relative stability of myelin in the mature dam.

**Fig 4.**
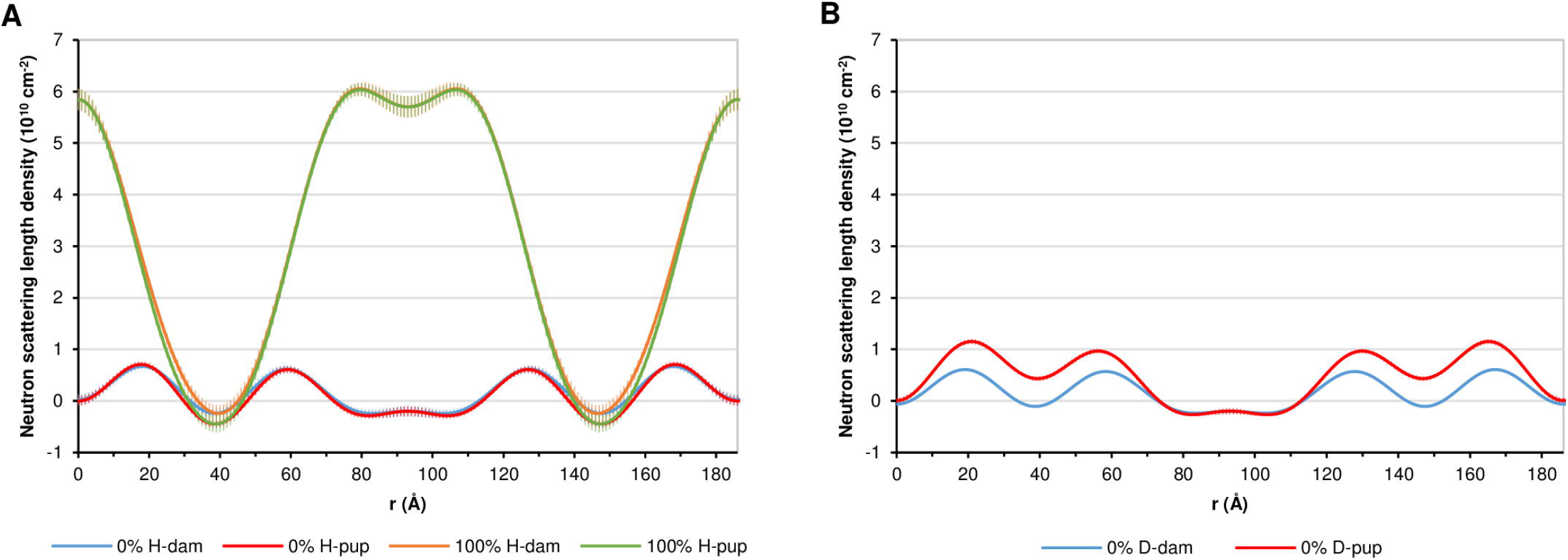
Scattering density profiles of myelin from pups and dams. The profiles were calculated using the interpolated structure amplitudes (for *h*=1–5) based on the linear regression shown in Fig 3. The origin (r=0 Å) corresponds to the cytoplasmic boundary between membrane pairs. The profiles were scaled such that in 0% and 100% D_2_O, the absolute density at the extracellular apposition was equal to the neutron scattering densities of H_2_O and D_2_O containing volume fractions of protein of 0.14, and in the center of the bilayer, a volume fraction of protein of 0.09 [7]. (A) H-dam vs. H-pup at 0% and at 100% D_2_O. (Profiles for H-pup in 0% and 100% D_2_O-PBS calculated from Bragg orders 1–6 are shown in S8 Fig.) (B) D-pup vs. D-dam at 0% D_2_O. (Data for D-dam at 100% D_2_O-PBS was not collected owing to time constraints on the beamline.)

Comparison of the scattering density profiles for myelinated nerves from the undeuterated vs. deuterated mice (H-mice vs. D-mice) revealed more clearly the localization of deuterium (Fig 5). Overall, the two profiles for H-dam and D-dam in 0% D_2_O-PBS were very similar but not identical (Fig 5A), as there was a slight but significant increase in scattering density for D-dam in the central hydrocarbon layer (Fig 5B; see also S9 Fig). This suggests that a smaller amount of deuterium had been incorporated into the dam’s tissue, most likely owing to the steady-state turnover of membrane constituents during the metabolism that maintains myelin’s integrity after maturation. By contrast with the small difference in scattering density profiles between the dams, the difference between the undeuterated and deuterated pups was characterized by a substantial elevation of scattering density centered in the membrane bilayer (Figs 5C,D and S7 Fig). This singular feature was evident and uniform across the entire range of contrast variation provided by different % D_2_O-PBS solutions (S6 and S7 Figs). Given that lipids are the major hydrogen-rich, non-aqueous constituent of myelin, constituting 70–80% of its dry weight [44], it is not altogether surprising that metabolically-incorporated deuterium would be most apparent in the H-rich hydrocarbon region of the membrane bilayer. Lipid deuteration would arise from incorporation of deuterium during *de novo* lipogenesis of deuterium from the D_2_O-spiked drinking water consumed by the pregnant dam and subsequently by the weaned pups. Studies on the incorporation of deuterium into fats (FA and TG) show that such labeling occurs directly from water, and also via enzyme-catalyzed H-D exchanges during metabolic conversions [45,46]. We expect that the myelin proteins are also deuterated; however, not only do they constitute only 20–30% of the myelin, but also they are distributed across the entire breadth of the membrane, which includes the intermembrane, polar spaces at the cytoplasmic and extracellular interfaces, and also the nonpolar transmembrane region. It has been estimated that hydrogens are ∼50% of all atoms in proteins, whereas they are >60% of all atoms in lipids (and closer to 70% in the hydrocarbon tails). Given the lower abundance of proteins compared to lipids in myelin, and their broader distribution across the membrane, our ability to detect—by neutron diffraction and with the current experimental design— deuterated protein in the presence of deuterated lipid would be challenging.

**Fig 5.**
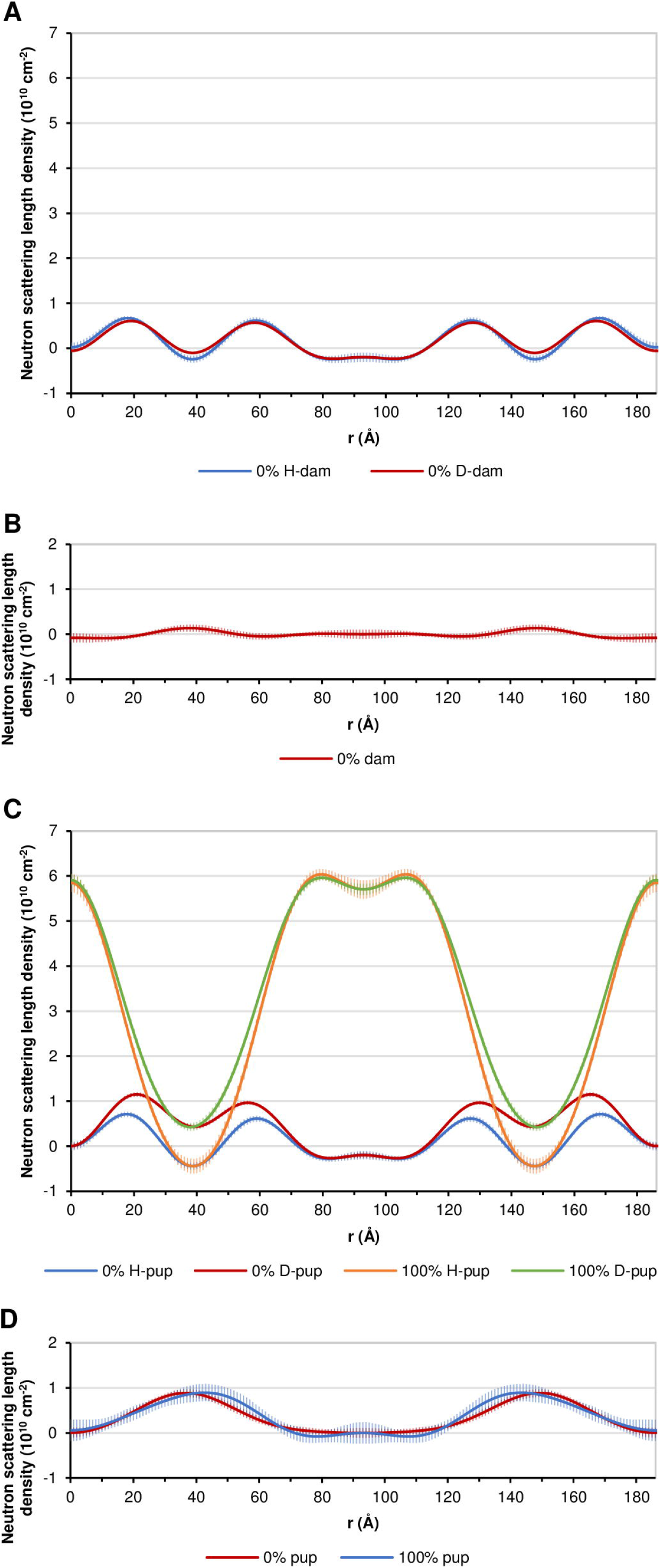
Scattering density profiles and difference profiles of myelin from D-mice vs. H-mice. The profiles were calculated using the interpolated structure amplitudes (for h=1–5) based on the linear regression shown in Fig 3. The origin (r=0 Å) corresponds to the cytoplasmic boundary between membrane pairs. The profiles were scaled as described in Fig 4 and text, and are all on the same scale. (A) Scattering density profiles for D-dam and H-dam in 0% D_2_O-PBS and (B) their difference profile. (For an expanded y-scale, see S9 Fig). (C) Scattering density profiles for D-pup and H-pup in 0% and 100% D_2_O-PBS, and (D) their difference profiles in 0% (red) and 100% (blue) D_2_O-PBS. (For an expanded y-scale for 0% D_2_O-PBS, see S9 Fig.) Comparison between D-pup and H-pup in the series of solutions of varying contrast showed this consistent difference (S6 and S7 Figs). Error bars for the difference profile (Δ = D ≈ H) were calculated from: □Δ = [(□D)^2^ + (□H)2]^½^, where □D and □H are the errors at each position in the neutron scattering density profile.

In addition to the prominent increase in scattering density in the hydrocarbon region, the difference maps across the range of contrast variation (S7 Fig) show a more subtle difference in deuterium distribution. As the amount of diffusible deuterium increased via H-D exchange from the equilibration solutions that ranged from 0% D_2_O to 100% D_2_O, there was a systematic shift of the high density, metabolically-incorporated deuterium-label peak by 7–8 □along the x-axis toward the lipid polar group layer in the extracellular half of the bilayer. At the current resolution of the data, this change is within the uncertainty of the error bars; however, it suggests that there are exchangeable hydrogens at this depth in the membrane bilayer, at least for aldehyde-fixed myelin.

We conclude from the structural analysis that the metabolically-incorporated deuterium is predominantly concentrated in the central trough of the membrane bilayer, i.e., where there is the greatest proportion of hydrogen atoms that could be substituted for in part by deuterium during *de novo* lipogenesis or during normal lipid turnover. To confirm the incorporation of deuterium into the myelin, and to gain a more thorough understanding of the nature of the deuteration, we undertook analysis by mass spectrometry of the lipids isolated from the trigeminal nerves that had been frozen in parallel to the sciatic nerves used for structural analysis.

### Metabolic deuteration of mouse tissue lipids

The earliest research describing the distribution of mass isotopologues among deuterated myelin lipids isolated from D_2_O-administered mice was reported by [13]. Their protocol was a “pulse-chase” in which 30% D_2_O in drinking water was available *ad libitum* for 4–6 weeks to four different groups of mice, ranging in age from five weeks to ∼2 years. The mice were sacrificed periodically for up to ∼200 days to follow the incorporation and disappearance of deuterium from myelin lipids purified from the CNS (cerebellum). An earlier study [47] focused on the replacement rates of liver lipids (TAG, PC, and PE) in ∼6-week old mice in which deuterium from D-spiked drinking water was integrated into fatty acids during a 15-day timespan. (We deduced the age of these mice from the stated, initial weight of the mice for the particular strain used, C3H/He, and its growth curve: https://www.criver.com/products-services/find-model/c3h-mouse?region=3611.) Both of these studies demonstrate the feasibility of incorporating deuterium—a stable non-radioactive isotope—to study lipid turnover in mice; however, all the mice including the youngest ones were beyond the age of their most rapid and robust myelination, which is up to ∼4 weeks in the PNS and ∼5 weeks in the CNS [48]. The idea to use MS to analyze isotopologue patterns in the biosynthesis of molecules in the fetus of a pregnant dam exposed to D_2_O-spiked drinking water was first mentioned in a review article in 2004 [49], but to our knowledge this experiment was not undertaken. To distinguish lipogenesis during myelination vs. that from myelin maintenance, we initiated D_2_O administration via drinking water to pregnant dams, and analyzed the lipids from both the dam and her pups when the latter were mature (P55). In the current paper, we report our MS analysis of trigeminal nerve lipids from the same mice in order to confirm and illuminate chemically our localization by neutron diffraction of the deuterium label that was centered in the membrane bilayer of sciatic nerve myelin.

### MS confirmed deuteration of lipids, and variable deuterium levels and label distribution in D-dam

Negative-ion MS of the lipids isolated from whole trigeminal nerves from the controls—H-dams and their offspring, H-pups—showed the expected natural abundance isotopes (^1^H, ^12^C, and ^14^N) for the monoisotopes (see upper two spectra in each panel of Fig 6, and upper spectrum of S11B Fig). As a control, we also used the MassWorks analytical tool to determine the level of naturally-occuring deuterium (^2^H, D) incorporation in the controls: the value was 0.19% ± 2.4% (n=230), which is about ten times the abundance of ^2^H in water [50]. Given the large standard deviation, this apparent low level of deuterium found in the controls is statistically indistinguishable from the expected zero value, which indicates MS measurement error or noise. By contrast with the MS spectra from the H-mice, all of the spectra from the D-pups and most spectra from the D-dam showed a decrease in the relative intensity of the monoisotopic peak that was paralleled by an increased number of detectable mass isotopologues (Fig 6, lower two spectra in each panel; and lower spectrum of S11B Fig). The increase in abundance of the evenly-spaced isotopologues at *m/z* values (A+1, A+2, …A+n) following the monoisotope peak (A_0_) indicated the addition of *N* = 1…..*n* deuterium atoms to the lipid, or the formatioon of M_1_, M_2_, …, M_n_ isotopologues related to the original undeuterated isotopologue M_0_. The MassWorks analysis clearly resolved this undeuterated isotopologue M_0_ from the deuterated isotopologues (M_1_, M_2_, …, M_n_) and indicated how much more deuterated the pups’ lipids were by contrast with the variable level of deuteration of the dam’s lipids: e.g., for the identified lipids, the average level of deuteration in the D-pups’ lipids was 97.6% ± 2.0% (n=54; range 89.6–99.6%), whereas their mother’s (D-dam’s) values ranged broadly from 11.4–97.3% (avg. 60.6% ± 26.4%, n=27) (Fig 7). Moreover, the number of additional isotopologues was more variable and generally fewer in D-dam. In other words, the percentage of lipids that had one or more deuterium atoms was substantially greater in the pups than in their mother. The consistent, overwhelming deuteration of lipids in the D-pups must be a consequence of the prolonged period from early fetal life to two months of age, when deuterium was incorporated metabolically into the rapidly myelinating tissues [48] and when there would likely be very slow turnover of lipids in the growing, multilamellar myelin sheath. During this same timeframe for treatment, the mother would have maintained a normal level of lipid turnover, which would depend not only on the particular lipid species but also on the molecular subspecies. This interpretation assumes that any effect of deuterium on biosynthesis and assembly, and on metabolism and turnover is negligible, notwithstanding that the C-D bond has a higher dissociation energy (341.4 kJ/mol vs. 338.4 for kJ/mol for C-H [51]) and is reported to be less susceptible to enzymatic cleavage [52]. Our finding for mouse PNS myelin is consistent with results from analysis of mouse and rat CNS myelin, which have been shown to undergo appreciable changes not only across phospholipid and glycolipid species but also among their molecular subspecies during development and maturation [53, 54].

**Fig 6.**
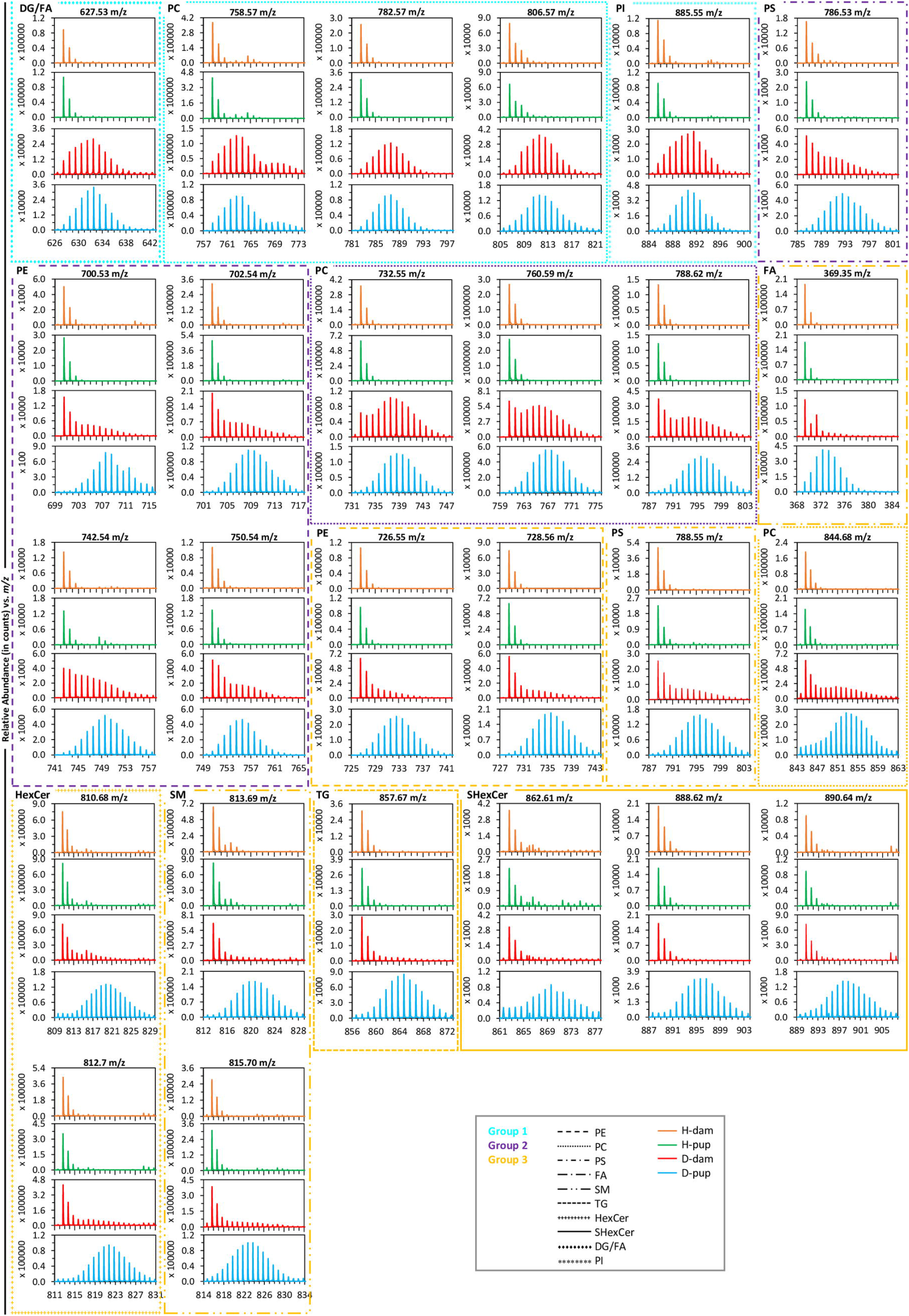
Mass spectrometry of lipids isolated from mouse trigeminal nerves. Each vertical set of four spectra (or “panel”) corresponds to a different lipid, and shows (top to bottom) the mass isotope distribution as relative abundance vs. *m/z* for H-dam, H-pup, D-dam, and D-pup. For undeuterated samples, the peaks correspond to natural abundance isotopes; and for deuterated samples, the series of peaks at increasing *m/z* values correspond to increasing incorporation of D. The panels are grouped by the extent of deuteration for D-dam: Group 1, high (bounded by blue); Group 2, intermediate (bounded by purple); and Group 3, low (bounded by bilious or olive green). Within each group of panels, lipids of the same species but different values of *m/z* are enclosed by a box that is distinguished by a particular line type (see key) and are identified in the upper left corner of the box by an abbreviation.

**Fig 7.**
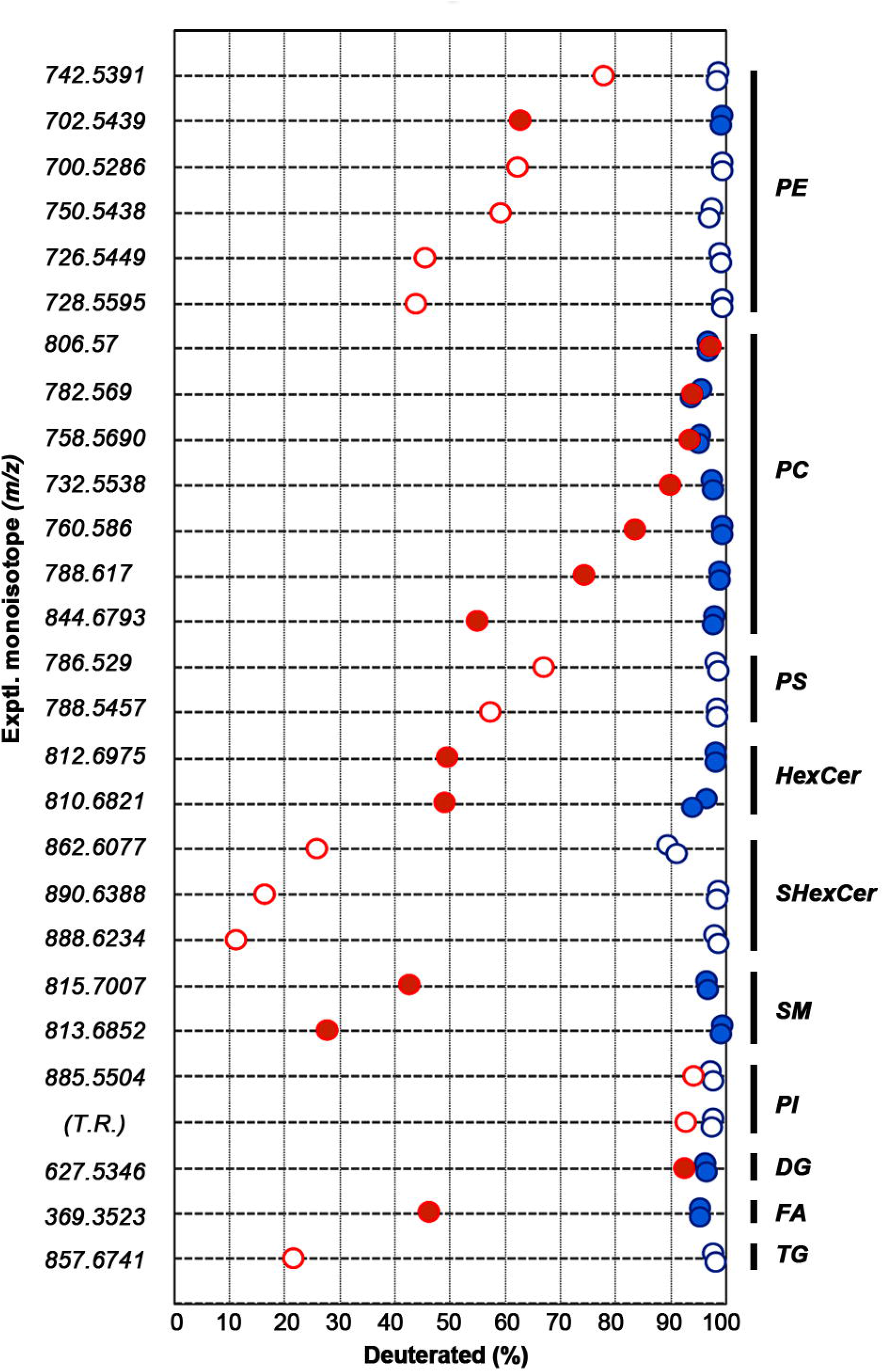
Comparison of deuteration levels of lipids in D-dam and D-pups. These results are based on mass spectrometry of lipids in negative and positive ion modes, as analyzed using MassWorks (Cerno Bioscience). Of the 89 experimental monoisotopes that we determined using MS-LAMP [20], 27 (y-axis) corresponded to *m/z* values that have been identified as specific lipid species in mouse or rat neural tissue and in curated databases (see references cited in Materials and methods). For each monoisotope value, the percent deuteration levels—calculated by summing the relative percentages of the deuterium isotopologues—are indicated for the mother (in red; D-dam) and two of her pups (in blue; D-pups). Seven different species of lipids plus three different species of acylglycerols were identified that corresponded to the *m/z* values: PE (6 sets of data), PC (7 sets), PS (2 sets), HexCer (2 sets), SHexCer (3 sets), SM (2 sets), and PI (1 set, and one technical replicate); and acylglycerols TG (1 set), DG (1 set) and FA (1 set). Open circles and filled circles correspond to measurements from negative and positive ion mode spectra, respectively. T.R. denotes technical replicate (see also Fig 8).

Based on the MS analysis of neural tissues from mice (and rats) and from curated databases, we identified 26 distinct lipids distributed over three different lipid classes (Fig 6) in our samples. The classes consisted of glycerophospholipids (with the species PE, PC, PS, and PI), sphingolipids (species HexCer, SHexCer, SM), and acylglycerols (species FA, DG, and TAG). Except for PI and the acylglycerols, each lipid species had more than one molecular subspecies. To compare lipids, we arranged the MS spectra by molecular subspecies according to the deuterium distribution patterns exhibited by D-dam, such that lipids with similar patterns were grouped together into three categories based on high (Group 1), intermediate (Group 2), and low levels of deuteration (Group 3), indicative of high, intermediate, and low turnover of lipids. The most heavily deuterated lipids (>90%) were PC (three of seven subspecies), PI, and DG, and closely resembled the corresponding spectra of the pups. The intermediate levels of deuteration (∼60–90%) also resembled that in the pups but at lower levels, and consisted of PE (four of six subspecies), PC (three of seven subspecies), and PS (one species). The least deuterated (∼10 to <60%), and hence most stable lipids, were SHexCer (three subspecies), TG, SM (two), PE (two of six), FA, HexCer (two), PC (one of seven), and PS (one of two).

Our results are consistent with previous findings that used precursors labeled with a variety of tracers (^13^C, ^14^C, ^2^H, ^3^H, ^32^P) and demonstrated, at least in the CNS, a differential distribution of label and widely differing turnover rates among the lipid classes of myelin. For example, there is general agreement that the ranking of mouse myelin lipids from most rapid to slowest turnover half-life is: (PI, PC) ≈ PE < (HexCer, SHexCer) < SM [13,55,56 and references therein].

What is distinctive about our results compared to the earlier studies is the clear variation in deuteration levels among the molecular subspecies of the lipids we identified: e.g., the seven subspecies of PC varied from ∼50% to >90%, and the six molecular subspecies of PE varied from ∼40% to ∼80%, indicating a wide range of metabolic turnover. In addition, all three subspecies of SHexCer had very low levels of deuteration (varying form ∼10% to <30%), reflecting long half-lives (stability) in the myelin membrane, whereas HexCer (the precursor for sulfatide) showed ∼50% deuteration and, therefore, greater turnover. The similarly-high deuteration levels (at >90%) of PI and DG is consistent with their role in the inositol-phospholipid signaling pathway, where hydrolysis of the membrane-bound, doubly-phosphorylated PI (or PIP2) forms two second messengers, one of which (DG) remains in the membrane. The deuterated fatty acyl chains of PI, therefore, most likely account for the deuterated fatty acyl chains of DG. The low level of deuteration for TG (∼20%) is consistent with its role as a stable, storage reservoir for fatty acids. The high deuteration level for several of the PC subspecies that we identified may relate to the biosynthetic pathway involving condensation between highly-deuterated DG and cytidine 5’-diphosphocholine (CDP-choline or citicoline) [57]. PC also plays a role in membrane-mediated cell signaling and phosphatidylcholine-transfer protein activation of other enzymes [58]. Taken together, our findings underscore not only the dynamic metabolic diversity of the lipids but also are consistent with widely varying functions among their subspecies.

As described above, deuterium enrichment was indicated by the progressive increase in the abundance of higher mass isotopologues. The distributions of these D-labeled molecules always formed symmetric, bell-shaped envelopes for the lipids of D-pups, but usually formed more complex envelopes for lipids of the D-dam. The symmetric shape of the envelope for D-pups indicated a maximum level of deuteration for the D-pups’ lipids, which depends on the relative amount of deuterium and hydrogen in the D_2_O-spiked drinking water. Exceptions were SHexCer (*m/z* 862.61) and PC (*m/z* 844.68) which showed slightly higher proportions of lower-mass isotopologues (e.g., M_1_ and M_2_)—this might be explained by different “schedules’’ of lipogenesis for these two lipid subspecies during gestation: e.g., these particular molecular subspecies of sulfatide and phosphatidylcholine might be the earliest synthesized by the myelin-forming cells of the gestating pups and thus crucial to subsequent myelin formation. There is abundant evidence in the CNS that sulfatide has crucial roles in oligodendrocyte maturation, and subsequently in myelin formation and stability (reviewed by [59]); and sulfatide subspecies have been shown to have different developmental timetables during oligodendrocyte maturation and myelinogenesis as demonstrated using imaging mass spectrometry [60]. Other distinct distributions were PE (*m/z* 700.53) and PC (*m/z* 758.57) which showed enhanced values for higher-mass isotopologues M_i_ (e.g., *i* > 10), suggesting a different population of these subspecies; however, these variations may also be accounted for by slight differences in the retention times (S11 Fig).

The average number of deuterium labels in a lipid, calculated as a weighted average, depended on the lipid species (Fig 8). By contrast with results from D-pups, the shapes of the isotopologue distribution curves for D-dam ranged from being symmetric, bell-shaped—Group 1 of D-dam (DG, PI, and PCs having *m/z* values of 806.57, 885.55, and 758.57 [the lattermost showing two, overlapping bell-shaped patterns])—to showing a complex combination of high-percentage monoisotope with nascent bell-shaped isotopologue distributions (many in Group 2), to being predominantly undeuterated with relatively small levels of label (Group 3). Similar isotopologue distributions were also reported for cerebellar myelin lipids [13] and for liver fatty acids [14,47] from mice given D_2_O-spiked drinking water. These researchers accounted for their findings by positing the coexistence of several fractions of molecules that included newly-synthesized ones having relatively few deuteriums, to molecules that were either undeuterated or synthesized from recycled deuterium-containing constituents. Reactions that are likely to be involved in incorporating deuterium from D_2_O into the lipids include glyceroneogenesis or glycolysis, where deuterium can be transferred into the C-H bonds of tryglceride’s glycerol backbone [45], and the fatty acid synthesis reaction, where deuterium can be incorporated enzymatically into the lengthening fatty acyl chains [46,61].

**Fig 8.**
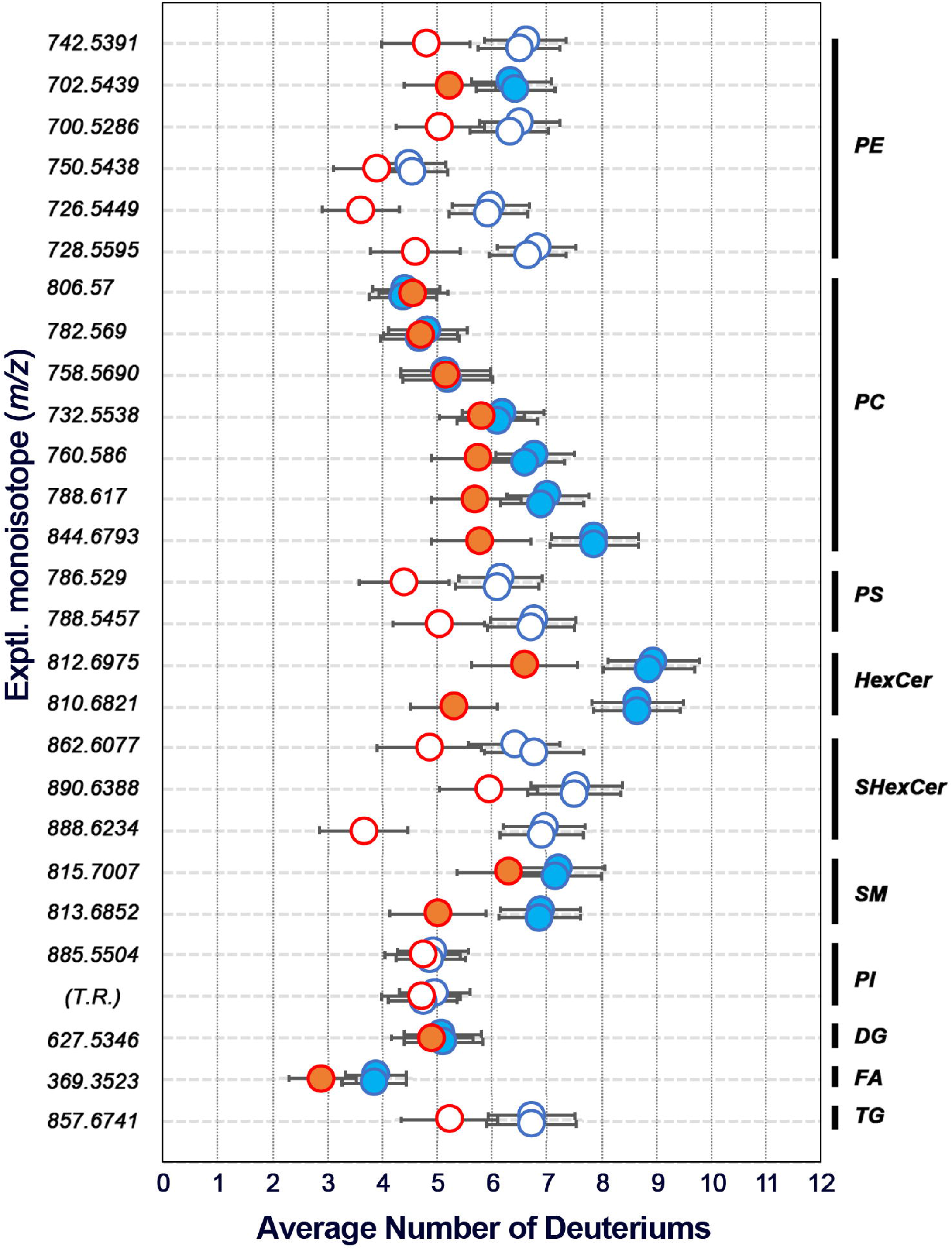
Average number of deuteriums per lipid species in D-dam and D-pups. The average number of deuteriums for each lipid species was determined by calculating the weighted average of the numbers of deuteriums incorporated for the corresponding D-dam (red) and D-pups (blue). Open and filled circles correspond, respectively, to measurements from negative and positive ion mode spectra. The error bars are standard errors calculated from the weighted standard deviations [36].

## Conclusions

Our study’s accomplishments were twofold: one, we demonstrated that neutron diffraction could be used to successfully localize in intact tissue the deuterium that had been metabolically incorporated *in vivo*; and two, we demonstrated that metabolic incorporation of deuterium as a stable isotope during gestation vs. after maturation can provide insight about the fates of specific lipid species and subspecies in myelin formation, maintenance, and turnover. Our experimental design—in which the pups were sacrificed at P55, which is immediately after the most rapid rate of myelination, whereas the dams were sacrificed at P131, near the end of the slowed rate of myelination [48]—enabled us to distinguish the different deuterium incorporation scenarios for *de novo* lipogenesis during myelin accretion vs. targeted turnover of lipids based on particular functional demands of mature myelin. Based on these findings, we anticipate exploitation of this approach to developing a comprehensive understanding of how lipids are differentially regulated throughout life—gestation, the three phases of myelin accumulation, myelin “stability” during its maintenance, and myelin aging—in normal and pathological tissue.

### Future perspectives

Our approach provides proof-of-concept for further research that relates specifically to myelin structure and dynamics and more broadly in other areas. In the experimental paradigm we used here for stable isotope incorporation via D_2_O-spiked drinking water, not only was myelin labeled, but a priori one can surmise that all tissues in the mice were deuterated—differentially for the pups vs. the dam. Analysis by neutron diffraction to localize metabolically-incorporated deuterium could be applied to tissues that exhibit macro- and supramolecular order, including muscle fibers, retinal rod outer segments, collagen fibers, keratin, bone, etc., and to proteins that could be purified and organized with crystalline or paracrystalline order, e.g., myelin proteins [62,63], hemoglobin, numerous enzymes, β- amyloid, etc. Recovering or preparing these different types of samples at different times during *in vivo* deuteration or after the D_2_O has been deleted from the drinking water, and analyzing by MS the changes in distribution patterns of deuterium labeling would illuminate our understanding of the dynamics of these tissues and molecular assemblies. In particular, with respect to myelin, we can ask the following questions: What functions account for the differential deuteration of PC subspecies in the dam? How do the turnover rates for the different lipids vary with age once the myelin is “fully” mature? When the balance in myelin formation and maintenance are upset owing to mutations in lipid biogenic enzymes or in myelin structural proteins, how are the lipid species’ turnovers affected? If lipids in the extracellular leaflet are less subject to turnover and more stable metabolically, can we argue that the <u>least</u>-deuterated lipids that we identified in D-dam are in the extracellular half of the bilayer—e.g., SHexCer, TG, and SM? Conversely, might we also argue that the <u>most highly</u>-deuterated lipids in D-dam are in the more metabolically-accessible cytoplasmic half of the bilayer—e.g., PC (three subspecies), PI, and DG? Thus, the convergence of lipidomics and metabolomics can be anticipated to provide extraordinary insight into myelinomics. An indication of the analytical and quantitative power of this approach has been demonstrated using HeLa cells [64,65]. In short, we anticipate that our correlated structural and biochemical study will promote a broad, comprehensive approach that is informative about structure, function, and metabolism in both health and disease.

## Supporting information

**S1 Fig. Feeding protocol for deuterium incorporation. S2 Fig. Determination of *d* from q**_**h**_ **vs. *h***.

**S3 Fig. Step-function model for fixed, metabolically-undeuterated myelin in D**_**2**_**O-PBS and H**_**2**_**O-PBS**.

**S4 Fig. Neutron diffraction patterns from sciatic nerves from metabolically-undeuterated (control) mice**.

**S5 Fig. Scattering density profiles for fixed vs. unfixed mouse sciatic nerve**.

**S6 Fig. Scattering density profiles for H-mice (*A and C*) and D-mice (*B and D*) and dams (*A and B*) and pups (*C and D*) for a series of solutions ranging from 0%–100% D**_**2**_**O-PBS**.

**S7 Fig. Localization of metabolically-incorporated deuterium**.

**S8 Fig. Scattering density profiles for H-pups in 0% (red) and 100% (blue) D**_**2**_**O-PBS solutions**.

**S9 Fig. Difference profiles between D- and H-pups (blue) and D- and H-dams (red) in 0% D**_**2**_**O-PBS**.

**S10 Fig. Thin-layer chromatograms of the whole lipid isolates from trigeminal nerves**.

**S11 Fig. Extracted ion chromatogram (EIC) from negative mode analysis of the whole lipid isolates from trigeminal nerves and mass spectra for H-dam and D-dam**.

**S1 Table. Number of sciatic nerve samples for neutron diffraction measurements**.

