## Supplemental Tables and Figures for "Metabolically-Incorporated Deuterium in Myelin Localized by Neutron Diffraction and Identified by Mass Spectrometry"

**List of Supporting Information**

**S1 Fig. Feeding protocol for deuterium incorporation.**

**S2 Fig. Determination of d from $q_h$ vs. *h*.**

**S3 Fig. Step-function model for fixed, metabolically-undeuterated myelin in $D_2O$-PBS and $H_2O$-PBS.**

**S4 Fig. Neutron diffraction patterns from sciatic nerves from metabolically-undeuterated (control) mice.**

**S5 Fig. Scattering density profiles for fixed vs. unfixed mouse sciatic nerve.**

**S6 Fig. Scattering density profiles for H-mice (*A and C*) and D-mice (*B and D*) and dams (*A and B*) and pups (*C and D*) for a series of solutions ranging from 0%–100% $D_2O$-PBS.**

**S7 Fig. Localization of metabolically-incorporated deuterium.**

**S8 Fig. Scattering density profiles for H-pups in 0% (red) and 100% (blue) $D_2O$-PBS solutions.**

**S9 Fig. Difference profiles between D- and H-pups (blue) and D- and H-dams (red) in 0% $D_2O$-PBS.**

**S10 Fig. Thin-layer chromatograms of the whole lipid isolates from trigeminal nerves.**

**S11 Fig. Extracted ion chromatogram (EIC) from negative mode analysis of the whole lipid isolates from trigeminal nerves and mass spectra for H-dam and D-dam.**

**S1 Table. Number of sciatic nerve samples for neutron diffraction measurements.**

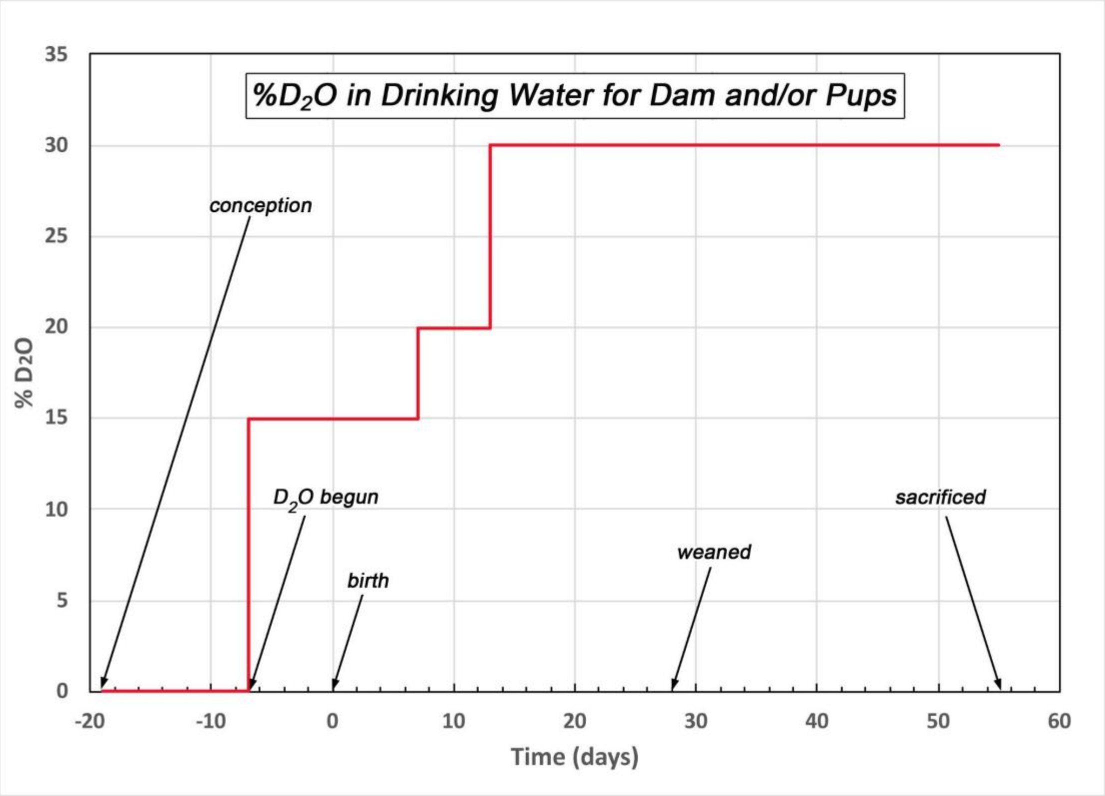


**S1 Fig. Feeding protocol for deuterium incorporation.** Pregnant C57BL/6 dams were initially provided ad libitum access to food and water. At gestational day 12, mice were separated into control (H-mice) and deuterium-treatment (D-mice) groups. For the latter, the drinking water was replaced with water containing 15% $D_2O$. At postnatal day 7, the volume fraction of $D_2O$ in the drinking water was increased to 20%; and at postnatal day 13, the $D_2O$ volume fraction in water was further increased to 30%. Pups were weaned at postnatal day 28, and both the dam and the pups were maintained on 30% $D_2O$-water. At postnatal day 55, all mice (D- and H-dams, D- and H-pups) were sacrificed and tissues were harvested for neutron diffraction and mass spectrometry.

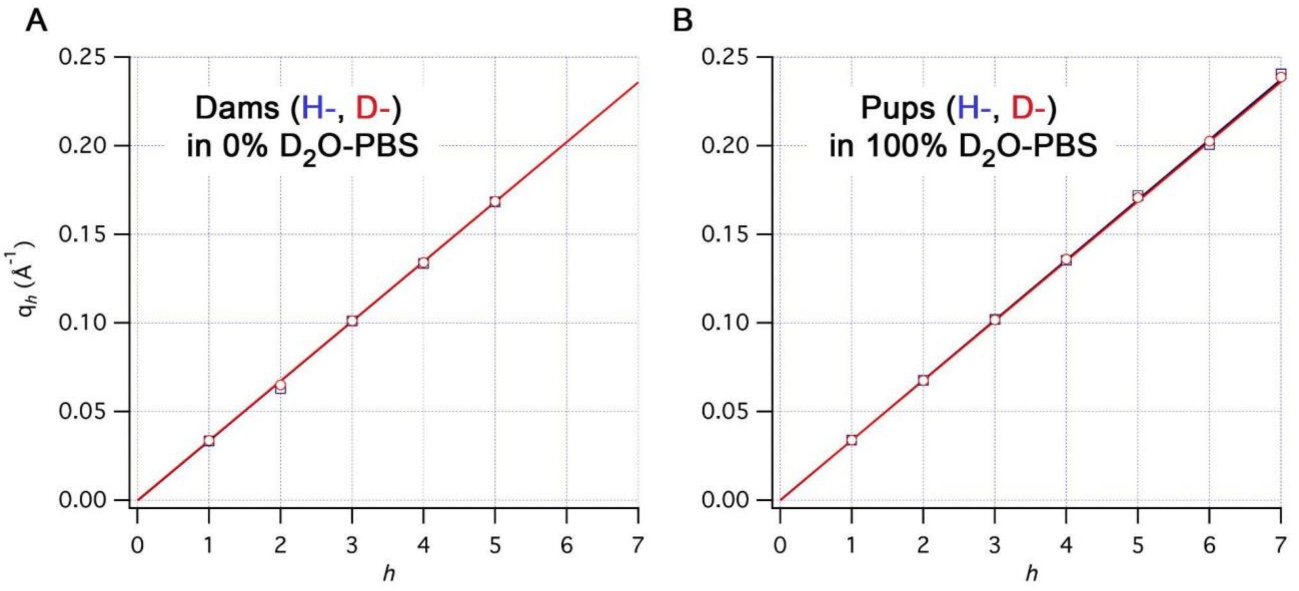


**S2 Fig. Determination of d from $q_h$ vs. h.** Myelin periodicity (*d*) measured from positions of Bragg peaks ($q_h$) as a function of Bragg order (*h*), for (A) H-dam (*blue*) and D-dam (*red*) in 0% $D_2O$-PBS, and for (B) H-pup (blue) and D-pup (red) in 100% $D_2O$-PBS. The period is given by $2\pi/b$, where *b* is the slope of the linear regression curve. The fixed sciatic nerves from the H- and D-mice, both dams and pups, showed virtually indistinguishable periods, indicating that the endogenous deuteration of the myelin had no effect on the width of the membrane pairs. The examples shown here are representative of the data across 17 samples, with 4 to 7 Bragg orders measured for each sample. The periodicity was 186.1 Å ± 0.1 Å. The measurements from the nerves are so coincident that it is difficult to see any differences between the H- and D- data points, with the exception of data for *h*=2 for the dams (A) and *h*=5–7 for the pups (B).

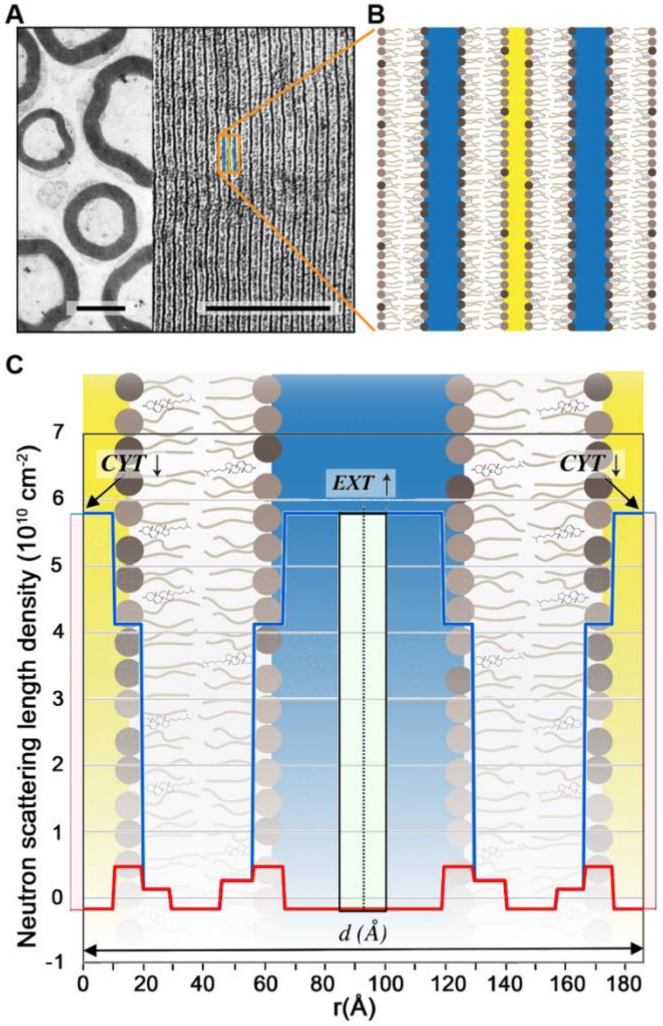


**S3 Fig. Step-function model for fixed, metabolically-undeuterated myelin in $D_2O$-PBS and $H_2O$-PBS.** The series of panels shows how the step-function model (C) is related to the ultrastructure of multilamellar myelin (A) and its lipid distribution (B). (A) Electron micrographs of a cross-section through a portion of a mouse sciatic nerve at low magnification to show the densely-stained, circular myelin sheaths (left; scale bar 3 μm) and at high magnification to show the periodic, paracrystalline stacking of the membrane pairs (right; scale bar 0.2 μm). (B) A schematic of the lipids in two different monolayers of polar heads (filled circles) and hydrocarbon tails (wavy lines), with cholesterol intercalated in the two monolayers. The bilayers, with their asymmetric composition of lipids, are arranged to simulate the symmetry of multilamellar myelin, where cytoplasmic faces are apposed across the narrow cytoplasmic space (yellow) and the extracellular faces are apposed across the extracellular space (blue). Specific lipids and myelin proteins, which include transmembrane and extrinsic polypeptides [1], have been omitted for clarity. (C) The neutron scattering length density (absolute scale [2]) is plotted vs. distance *r* (Å) from the origin at the cytoplasmic apposition. The center of the bilayer is at 37.1 Å [3]; and the myelin period for the fixed nerves is 186.1 Å (S2 Fig). In the fixed myelin (mouse sciatic) the cytoplasmic space (*CYT*) was narrower by 4–5 Å (pale pink areas) whereas the extracellular spaces (*EXT*) was wider by 15–16 Å (pale green areas) compared to unfixed tissue. The down and up arrows indicate a decrease or increase in the width of the space between membranes, as a consequence of chemical fixation. The blue and red indicate the step-models for myelin in $D_2O$- and $H_2O$-PBS, respectively. The schematic behind the scattering length density models shows how the lipid hydrocarbon tails, cholesterol's steroid nuclei, the lipid polar head groups, and the water-accessible regions at the extracellular and cytoplasmic define the step-function. These simple step-models were what we used to calculate continuous, membrane pair transforms which when sampled at *h/d*—where *h* is the Bragg order and the period *d*=186.1 Å—were used to predict the structure amplitudes *F(h/d)* (see Text, Fig 2).

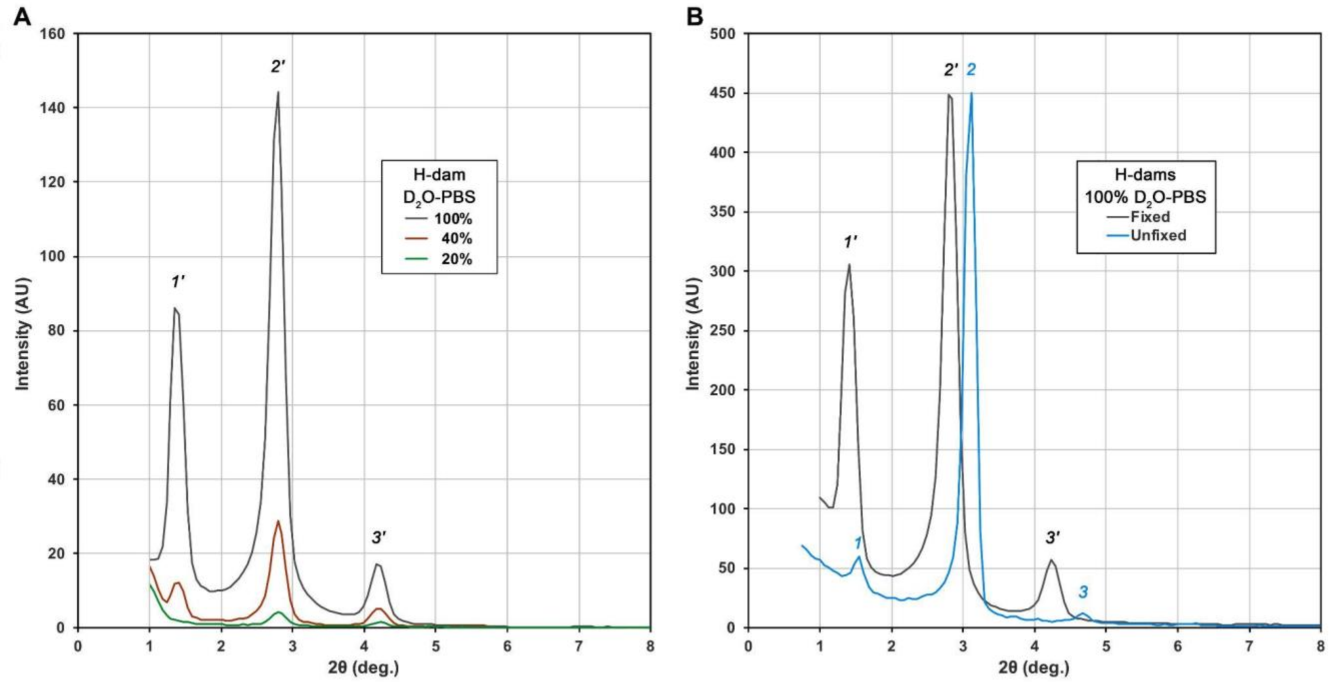


**S4 Fig. Neutron diffraction patterns from sciatic nerves from metabolically-undeuterated (control) mice.** (A) H-dam nerves equilibrated at 100%, 40%, and 20% $D_2O$-PBS. Only the first three Bragg orders are shown here, owing to the substantially weaker intensity of the higher orders (see Text, Fig 1). The intensities changed as $D_2O$ isomorphously replaced $H_2O$ in the water-accessible regions of the multilamellar membrane arrays. That the change in intensities with $D_2O$ was systematic is evident from a plot of the changes in the structure amplitudes measured from the intensities (see Text, and Figs 2 and 3). (B) Comparison of unfixed [4] and fixed (current paper) nerves in 100% $D_2O$-PBS. The intensities were scaled to the 2nd order reflection (at a scattering angle of ~3°). Primed and unprimed numerals denote the Bragg orders for the fixed and unfixed myelin, respectively. Note the change in intensities and shift in positions of the peaks to smaller scattering angle for the fixed myelin, indicating an increase in myelin periodicity and consistent with a change in membrane packing [3].

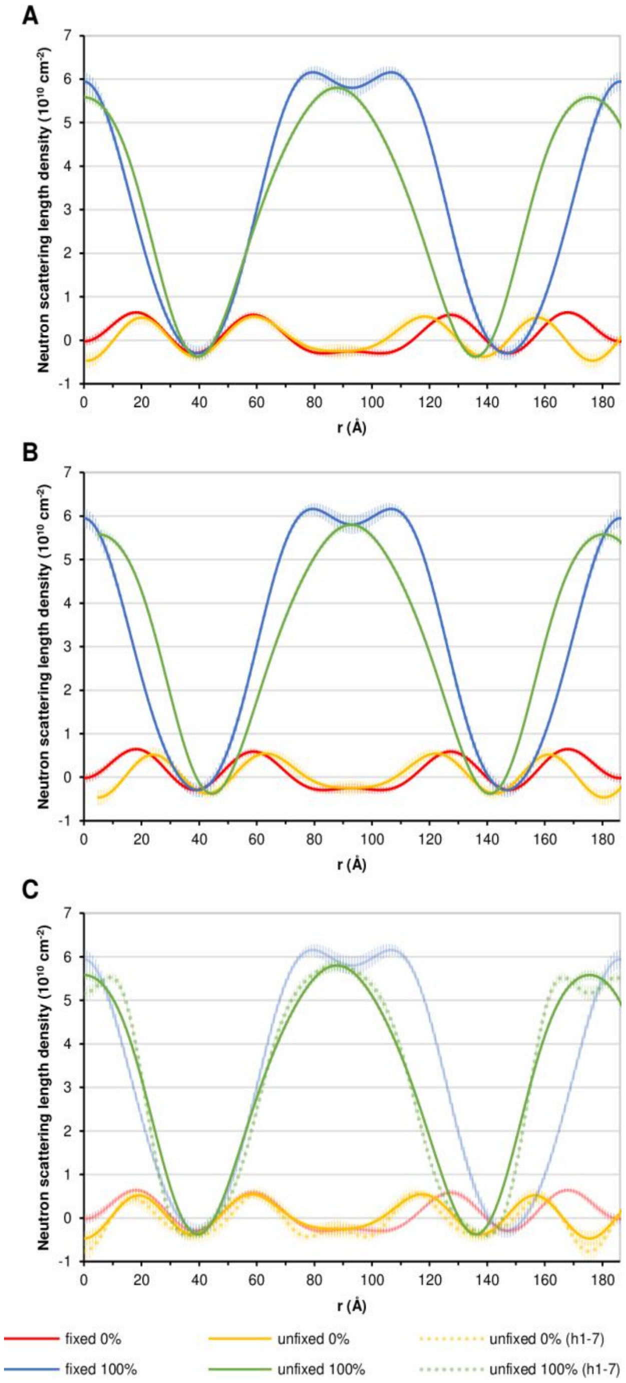


**S5 Fig. Scattering density profiles for fixed vs. unfixed mouse sciatic nerve.** The profiles for myelin in 0% and 100% $D_2O$-PBS were calculated for fixed myelin using the interpolated structure amplitudes (for $h$ = 1–5) based on the linear regression shown in Fig 3 and for unfixed myelin according to [4]. The origin ($r$=0 Å) corresponds to the cytoplasmic boundary between membrane pairs. The error bars (as short vertical bars) for the fixed samples were calculated from $\delta$F values (see Materials and methods), and for unfixed samples from counting statistics [4]. The neutron scattering density (y-axis) units are $10^{10}cm^{-2}$. (A) The structural similarity between the pair of profiles for unfixed (yellow) and fixed (red) myelin at 0% $D_2O$-PBS is striking, particularly at the hydrocarbon trough ($r$~40 Å) and the bounding lipid head groups, supporting the phasing of the reflections. The main difference is the slight elevation in scattering density at the cytoplasmic boundary and compaction of the cytoplasmic space. When the two membranes are compared at 100% $D_2O$-PBS, a different distribution of $D_2O$ is evident at the extracellular apposition, comprising the region from ~70–120 Å. (B) Plotting the profiles relative to the center of the extracellular boundary (at r~100 Å) shows more clearly the widening of this boundary in fixed myelin. The broader distribution of $D_2O$ is evident for fixed myelin in 100% $D_2O$-PBS (blue), as is the flatter boundary region for fixed myelin in 0% $D_2O$-PBS (red). (C) Including the higher Bragg orders ($h$ = 6 and 7) in calculating profiles for unfixed and fixed myelin in 0% and 100% $D_2O$-PBS resolves into two peaks the high scattering density of the cytoplasmic apposition for unfixed myelin in 100% $D_2O$-PBS (green, dotted vs. continuous curve).

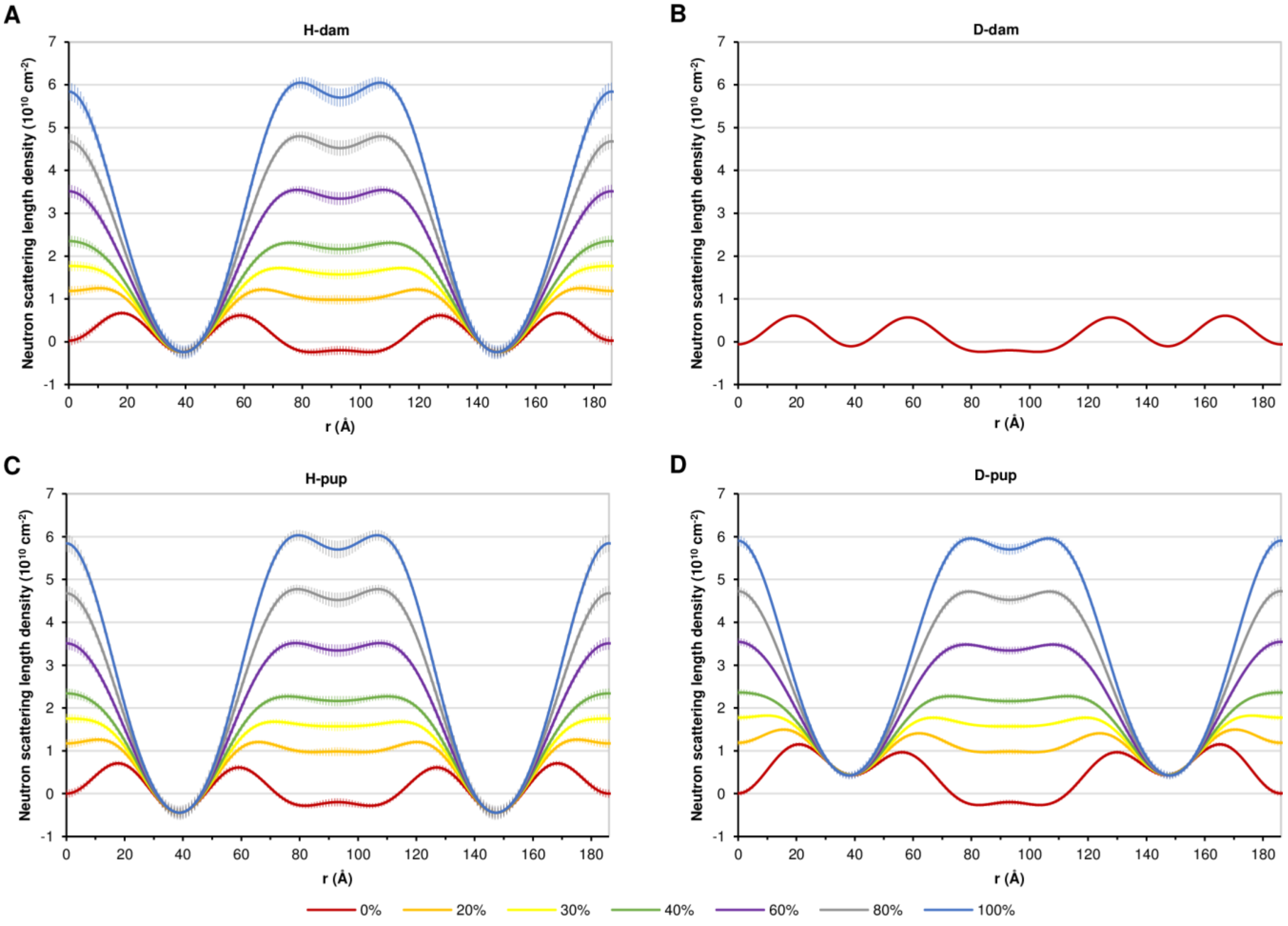


**S6 Fig. Scattering density profiles for H-mice (A and C) and D-mice (B and D) and dams (A and B) and pups (C and D) for a series of solutions ranging from 0%–100% $D_2O$-PBS.** Comparison of corresponding profiles for H-pup (C) vs. D-pup (D) having the same %$D_2O$-PBS shows that the major difference between these pairs of profiles is a consistent elevation of neutron scattering density centered in the lipid bilayer trough of the D-mice, at r ≈40 Å and r ≈145 Å. (The profiles for 0% and 100% $D_2O$-PBS are also shown in Fig 5 of the main text). A similar difference, though less obvious, is also observed for H-dam (A) vs. D-dam (B) at 0% $D_2O$-PBS.

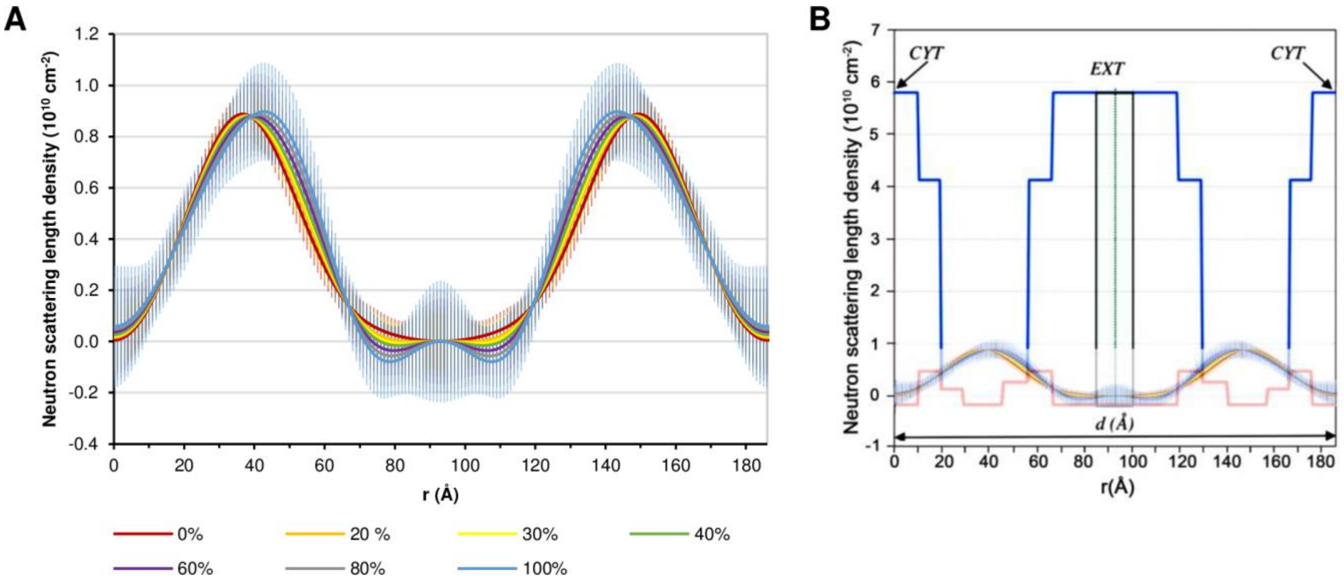


**S7 Fig. Localization of metabolically-incorporated deuterium.** (A) Difference profiles for H-pup and D-pup in a series of $D_2O$-PBS solutions ranging from 0%–100% $D_2O$-PBS, show a virtually uniform localization of deuterium across the different conditions. The difference profiles for 0% and 100% $D_2O$-PBS (red, blue), also shown in Fig 5D of the main text, are on an expanded y-axis here to show more clearly the systematic but apparently not-significant shift of the high-density peak toward the extracellular apposition at higher $D_2O$ content in the PBS. (B) Superposition of the difference profiles (A) onto the step-function model (S3 Fig) shows that the deuterium is predominantly concentrated in the hydrocarbon region of the membrane array.

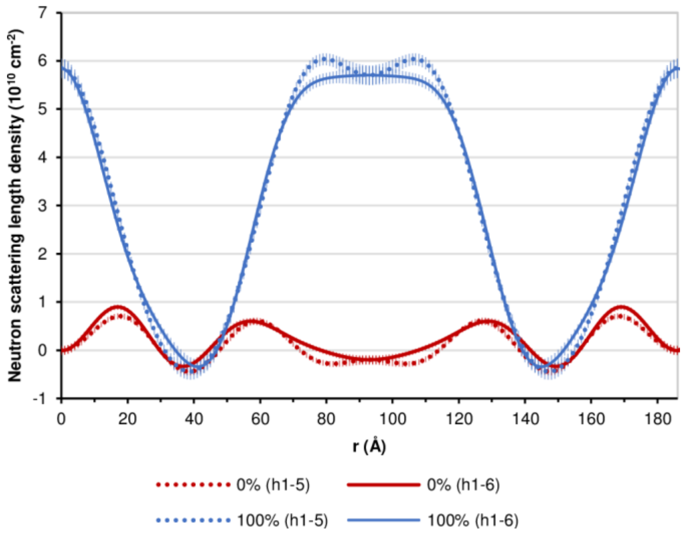


**S8 Fig. Scattering density profiles for H-pups in 0% (red) and 100% (blue) $D_2O$-PBS solutions.** The profiles were calculated for Bragg orders 1–5 (dotted curve) and orders 1–6 (solid curve), with phases assigned according to Table 1 (see also Figs 2C and 3F). Inclusion of the weak 6$^{th}$ order essentially flattens the neutron scattering density at the extracellular apposition.

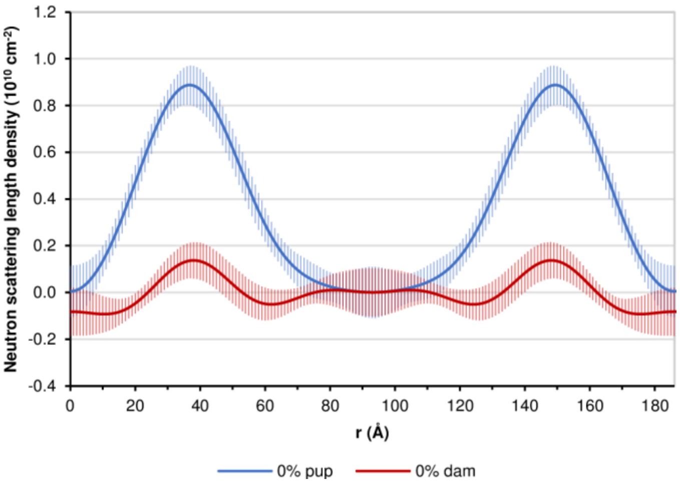


**S9 Fig. Difference profiles between D- and H-pups (blue) and D- and H-dams (red) in 0% $D_2O$-PBS.** The profiles are plotted at the same x-axis scale as the neutron density scattering profiles (Figs. 4 and 5), but on an expanded y-axis to show more clearly the slight, but significant increase in scattering density at the center of the hydrocarbon region (r~40 Å) for the dam.

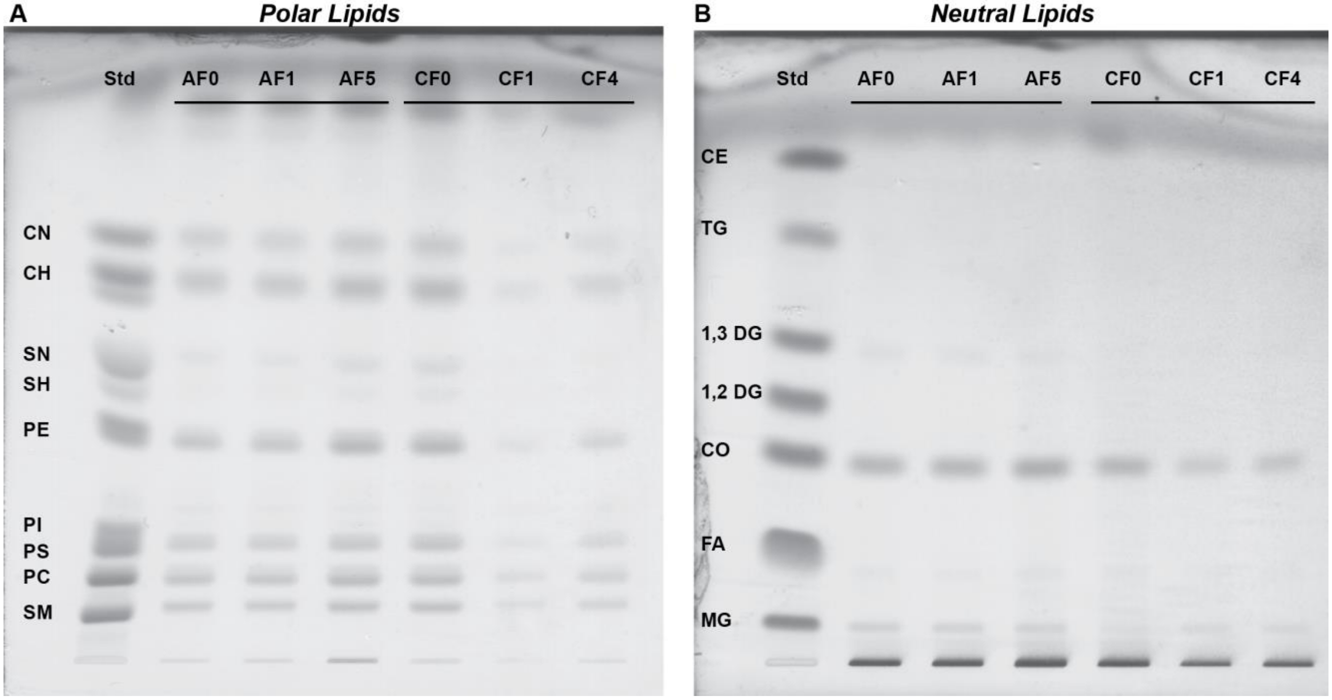


**S10 Fig. Thin-layer chromatograms of the whole lipid isolates from trigeminal nerves.** Non-deuterated ("AFn") and deuterated ("CFn") samples from the dams (control "AF0" and treated "CF0") and from ~55 day old, female pups (controls, "AF1", "AF5"; treated, "CF1", "CF4") were compared against standard lipids (Std). (NB. These "designations" are arbitrary sample coding names.) (A) Polar lipids: CN, non-hydroxy cerebroside; CH, hydroxy cerebroside; SN, non-hydroxy sulfatide; SH, hydroxy sulfatide; PE, phosphatidylethanolamine; PI, phosphatidylinositol; PS, phosphatidylserine; PC, phosphatidylcholine; SM, sphingomyelin. Note that in the Text, we use the abbreviations HexCer for cerebrosides and SHexCer for sulfatides, according to suggested standardized notation for mass spectrometric data [5]. (B) Neutral lipids: CE, cholesterol esters; TG, triglycerides; 1,3 DG and 1,2 DG, diglycerides; CO, cholesterol; FA, free fatty acids; MG, monoglyceride.

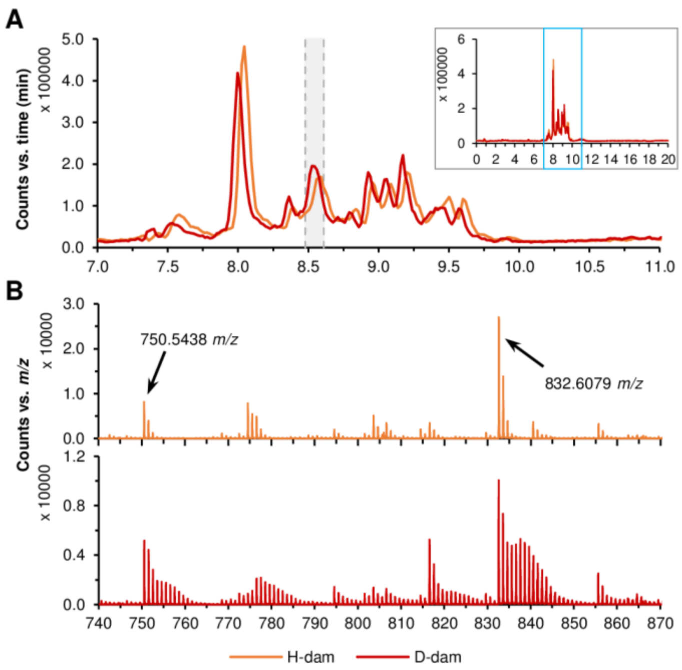


**S11 Fig. Total ion chromatogram (TIC) from negative mode analysis of the whole lipid isolates from trigeminal nerves and mass spectra for H-dam and D-dam.** (A) Close-up TIC of H-dam (orange) and D-dam (red) is shown as counts vs. time. Retention time window used for the mass spectra averaging is shown in gray. *Inset*: Whole TIC of H-dam and D-dam. The blue box shows the range of the close-up. The x- and y- axes have the same units as the close-up. (B) Spectrum of H-dam (upper panel) and D-dam (lower panel) of the peak at 8.48–8.61 min showing, for example, ions at *m/z* 750.5438 and 832.6079, which were identified as PE and PS, respectively, using LIPID MAPS [6,7]. Whereas PE was multiply-vetted as a mouse myelin lipid (see Table 2 for references), PS was not so vetted.

**S1 Table. Number of sciatic nerve samples for neutron diffraction measurements**.

| | $D_2O$ soaking | | | | | |
|---|---|---|---|---|---|---|
| | **100 %** | **60 %** | **40 %** | **30 %** | **20 %** | **0 %** |
| H-dam | 1 | | 1 | | 1 | 2 |
| H-pup | 3 | 3 | 2 | 2 | 3 | 6 |
| D-dam | | | | | | 2 |
| D-pup | 3 | 3 | 2 | 2 | 3 | 6 |
